# CAFE: A Co-folding Approach for Fragment Exploration of Allosteric and Cryptic Binding Sites

**DOI:** 10.64898/2026.07.19.739466

**Authors:** Justin Purnomo, Kunyang Sun, Teresa Head-Gordon

**Author notes:** Corresponding author, July 20, 2026.

## Abstract

Co-folding models hold immense potential for allosteric drug discovery, but have been severely hampered by their systematic bias toward orthosteric ligand binding. While fragment screening has been proposed for allosteric binding site discovery, we show that co-folding models still suffer from memorization in which chemically simpler fragments also default to canonical orthosteric binding sites. To overcome these limitations, we introduce CAFE (Co-folding Approach for Fragment Exploration), a co-folding protocol that uses competitive orthosteric blockers to divert fragments into non-canonical sites as illustrated here with the Boltz-2 co-folding model. Using ADP as an orthosteric blocker for the kinase family, we find CAFE substantially increases the allosteric binding site exploration for fragments, with notably strong absolute binding free energies that match or exceed those of known crystallographic poses, without post-hoc refinement of the Boltz-2 prediction. We also show that CAFE identifies cryptic binding pockets undetected by conventional pocket prediction tools, some of which are more thermodynamically favorable than the allosteric or orthosteric pockets. To demonstrate generality, we apply CAFE using Type I orthosteric blockers for kinase proteins, known orthosteric ligands as blockers for non-kinase proteins in the RAS-MAPK signaling pathway, and for virtual screening campaigns using fragment libraries for new fragments that selectively engage allosteric and cryptic binding sites. CAFE establishes orthosteric blocking and fragment screening as a training-free, inference-time protocol that helps overcome some of the limitations of current co-folding models while elevating their great promise for allosteric and cryptic binding drug discovery.

## 1 Introduction

Allosteric inhibition offers a powerful strategy to achieve target selectivity across highly homologous protein families in order to bypass orthosteric resistance mutations^1,2^. However, rational allosteric drug discovery remains notoriously difficult because allosteric pockets are structurally diverse, often cryptic or transient, and lack the conserved pharmacophoric features that guide drug discovery campaigns at orthosteric sites^3^. Traditionally, allostery has relied on expensive enhanced-sampling molecular dynamics^4–6^, but recent deep learning structure prediction tools offer a promising and new computational path forward to address allosteric and cryptic binding site discovery. Co-folding models, including AlphaFold3^7^, Boltz-1 and Boltz-2^8,9^, Chai-1^10^, RoseTTAFold All-Atom^11^, and related architectures^12–15^, jointly predict protein and complete ligand coordinates directly from sequence and molecular graph inputs, enabling pose prediction without a pre-existing receptor structure. These models achieve competitive performance on established benchmarks and have been widely adopted as first-pass docking alternatives^16,17^. For allosteric drug discovery in particular, co-folding models are attractive because they capture induced-fit effects implicitly through joint protein-ligand diffusion, allowing the model to sample pocket geometries that are accessible only in the presence of the ligand, a capability that rigid-receptor docking methods struggle to match^18,19^.

However, more recent benchmarks demonstrate that co-folding models systematically struggle with allosteric ligand pose prediction, primarily due to the sparse representation of allosteric complexes in the Protein Data Bank (PDB) and the difficulty of capturing large conformational transitions^20–22^. This failure is widely hypothesized to be protein-driven, reflecting learned model biases toward dominant orthosteric sites^23^. This problem is most acute in the kinase family, where the highly conserved adenosine triphosphate (ATP) orthosteric binding site acts as a potent structural attractor across hundreds of homologous enzymes. While allosteric kinase inhibitors—such as the myristoyl-pocket binder asciminib^24^ and the MEK inhibitor trametinib^25^—demonstrate the immense clinical value of escaping the ATP pocket to achieve target selectivity and bypass resistance mutations^1,2^, data-driven orthosteric memorization severely hinders their computational discovery using co-folding models.

To improve allosteric placement, Chen et al. fine-tuned co-folding diffusion models on a curated allosteric dataset^26^, but such approaches remain constrained by the sparse allosteric training data and fail to generalize to uncharacterized binding sites. Beyond explicit retraining and fine-tuning, other inference-time strategies include duplicating the allosteric ligand in the input to enhance sampling^27^, introducing known site occupants as extra chains to block competing orthosteric pockets^28^, or on deep-learning tools like AF2BIND and AlphaFold ensembles to predict binding residues and open states^29,30^. Crucially, however, these methods rely on whole, full-sized ligands bound to pre-characterized sites and do not address the upstream challenge of de novo allosteric site or cryptic site discovery in the absence of prior structural knowledge.

Fragment-based drug discovery (FBDD) offers a compelling strategy to address these gaps. Because chemical fragments sample structural space efficiently and can more readily bind to new pharmacophoric hot spots, they are uniquely equipped to map alternative regulatory sites^31^. Unfortunately, our study reveals that orthosteric memorization using co-folding models is still a problem even with fragments, such that even minimal probes containing only aromatic rings, hydrogen bond acceptors, and/or nitrogen atoms default to the orthosteric site regardless of their parent ligand’s origin or chemical context. However, experimental fragment screening has proven that fragments can identify novel allosteric pockets when orthosteric occupancy is used to direct fragment binding away from the active site, such as soaking GALK1 crystals pre-loaded with an active-site ligand^32^. Once favorable non-orthosteric sites are found using fragments, they can be elaborated into larger molecules for lead optimization by improving their drug-like profile.

Motivated by the experimental logic of active-site-occupied crystal soaking combined with fragment screening, we introduce CAFE (Co-folding Approach for Fragment Exploration), a simple and training-free inference protocol designed to break fragment memorization and discover allosteric and cryptic binding sites de novo. CAFE incorporates competitive orthosteric blockers directly into the co-folding input for Boltz-2, physically excluding fragments from canonical pockets during joint diffusion. We perform an extensive analysis across the kinase family, employing adenosine diphosphate (ADP) as a universal orthosteric blocker, as well as Type I kinase blockers, alongside a library of BRICS-derived fragments^33^. Across kinome targets, CAFE increases the mean rate of fragment co-folding to non-orthosteric sites from 11.7% up to 85.0%. Crucially, absolute binding free energy (ABFE) calculations confirm that CAFE-discovered fragment poses achieve thermodynamic binding affinities matching or exceeding crystallographic poses without requiring post-hoc structural refinement^34,35^. We also show that CAFE can build on the promise of co-folding models by identifying cryptic pockets undetected by special purpose pocket-searching software^36,37^, using co-folding with both orthosteric and allosteric blockers, and as illustrated by the discovery of a CK2α cryptic site that is more thermodynamically favorable for fragment binding than the canonical binding pockets. We demonstrate the broader applicability of CAFE by its extension to non-kinase targets illustrated by the phosphatase PTP1B and GTPase KRAS when appropriate orthosteric ligands are identified as blockers, and how CAFE can be used in a virtual screening campaign, where we are able to identity fragments from Enamine that selectively engage allosteric or cryptic binding sites. In summary, CAFE establishes orthosteric blocking and fragment screening as a training-free inference-time protocol that converts co-folding bias from a fundamental limitation into a powerful engine for AI-driven allosteric drug discovery.

## 2 Methods

Figure 1 illustrates the CAFE workflow. We first describe the selection of kinase and non-kinase targets and associated orthosteric and allosteric reference structures. This is followed by the approach to fragment library creation, an outline of the general co-folding blocking strategy, and metrics used to score fragment pocket localization. Finally we present the absolute binding free energy (ABFE) protocol used to assess the thermodynamic viability of predicted poses.

**Figure 1:**
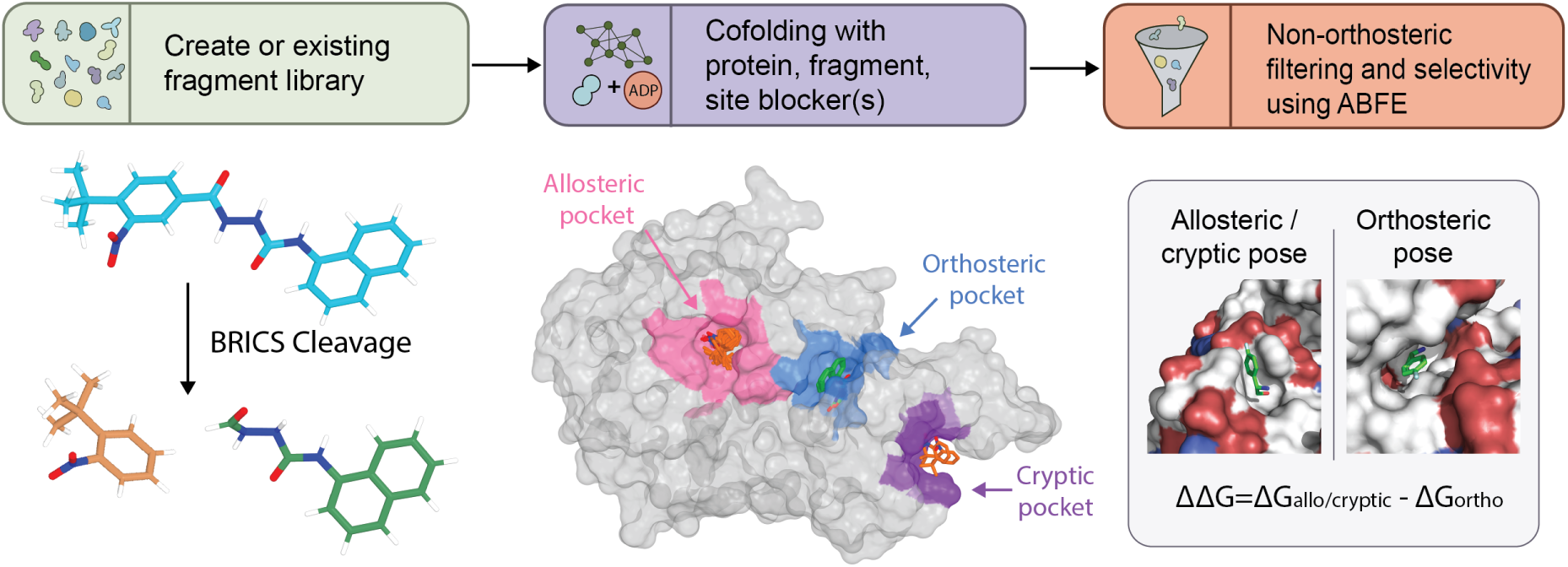
The CAFE prospective screening workflow and pocket-selective fragment recovery. In this work we use BRICS to create a fragment library from a whole ligand source such as Enamine fragments. Fragments are co-folded with each target protein under orthosteric blocking ligands, and then filtered for non-orthosteric localization. Absolute binding free energies (referenced to aqueous solvent) of non-orthosteric binding are compared to the free energies of co-folding without blocking, (ΔΔ*G* = Δ*G*_allo/cryptic_ − Δ*G*_ortho_) to confirm blocker-dependent selectivity.

### 2.1 Protein target and allosteric reference site selection

Five human kinases were selected for which multiple deposited allosteric inhibitor co-crystal structures are available: AKT2 (UniProt P31751), CDK2 (P24941), CHEK1 (O14757), CSNK2A1 (P68400), and MAPK14 (Q16539). Together these targets span a wide range of orthosteric structural representation in the PDB^38^, providing a deliberate test of whether ADP-blocked co-folding generalizes across targets with varying degrees of training data coverage.

For each kinase, one orthosteric reference structure was selected from independently deposited type-I inhibitor co-crystals: 2JDO (AKT2), 2UUE (CDK2), 2YEX (CHEK1), 3WAR (CSNK2A1), and 3S3I (MAPK14). These structures define the orthosteric center-of-mass (COM) used for localization scoring and serve as the source of the Type I blocker SMILES in the orthosteric blocker control arm. Allosteric reference structures were drawn from all available co-crystal structures with ligands bound at non-orthosteric sites for each UniProt entry: 8Q61, 9C1W (AKT2); 6Q3F, 6Q49, 6Q4K, 8VQ3, 8VQ4 (CDK2); 3F9N, 3JVR, 3JVS (CHEK1); 5MMF, 5MOD, 5OSU, 6GIH (CSNK2A1); and 3NEW, 4E6A, 4E6C, 4E8A, 5N63, 5N64, 5N67, 5N68, 8X3M, 8YD9 (MAPK14). Fragment localization to the allosteric region was scored against all available allosteric references simultaneously, taking the minimum COM distance across the full reference set. Pocket separation between orthosteric and allosteric reference COMs ranged from 12.8 Å (AKT2) to 38.0 Å (CDK2), confirming sufficient geometric separation for unambiguous localization scoring.

To assess whether the blocked co-folding framework extends beyond kinases, two non-kinase targets were also selected using target-appropriate orthosteric and allosteric reference structures. For PTP1B (UniProt P18031), the orthosteric reference was PDB 5K9W (catalytic cleft occupied by the phosphotyrosine-containing hexapeptide DADEpYL), with allosteric references drawn from seven structures spanning the BB-loop and *α*7 allosteric site (1T49, 1T4J, 7GSA, 7GTQ, 8G65, 8G68, 8G69). For KRAS (P01116), the orthosteric reference was PDB 4OBE (GDP; PubChem CID 8977), with allosteric references targeting the switch II pocket (7RPZ, 7U8H, 5V71).

### 2.2 Fragment library construction

The fragment library was derived from two cohorts of parent kinase ligands from the five kinases described above and decomposed by single-cut BRICS fragmentation^33^ using RDKit^39^. Orthosteric fragments were derived from 22 type-I co-crystal parents: 19 from a curated Modi–Dunbrack filtered type-I structure set (wild-type, non-phosphorylated, non-covalent holo inhibitor co-crystals)^40^, and three AKT2 structures (3E88, 3D0E, 2JDO) curated independently to ensure sufficient AKT2 representation. Allosteric fragments were derived from 21 of the 24 allosteric reference structures listed above; 4E6A, 4E6C, and 4E8A were excluded because deposited ligand coordinates represent incomplete lipid head groups relative to the catalog parent SMILES and could not support reliable native crystal fragment builds.

For each parent SMILES, all retrosynthetically labeled cleavable bonds were identified and cut independently, yielding two products per cleavage site. Dummy atoms at cleavage sites were capped with hydrogens, fragments with ≥ 8 heavy atoms and molecular weight ≥ 150 Da were retained, and fragments within each cohort were deduplicated on canonical isomeric SMILES. Parents lacking BRICS-cuttable bonds but satisfying the heavy-atom cutoff were retained as intact whole-ligand fragments (e.g., CDK2 HEW/HGQ). The complete set of fragment IDs and SMILES strings are presented in Supplementary Table S1. Chemical overlap between cohorts was assessed using Morgan fingerprints (radius = 2, 2048 bits)^41^. UMAP embedding of the 116 unique orthosteric and 116 unique allosteric fragments (Morgan fingerprints, Jaccard metric) confirmed that the two cohorts occupy distinct regions of chemical space with no cross-panel SMILES overlap (Supplementary Figure S1).

To test whether active-site blocking promotes non-orthosteric exploration outside kinases, we applied the single-cut BRICS procedure mentioned above to curated parent ligands for PTP1B and KRAS. For PTP1B, seven allosteric ligands were curated (1T49/892, 1T4J/FRJ, 7GSA/FM0260, 7GTQ/S7S, 8G65/DES4799, 8G65 and 8G69), yielding a final screen set of 23 fragments. For KRAS, three allosteric parents were curated (7RPZ/6IC, 7U8H/LX6, 5V7I/8ZG)), yielding 14 fragments.

As an independent chemical-space probe distinct from BRICS decomposition of co-crystal parents, we screened a diverse subset of the commercial Enamine High Fidelity Library (1,920 plated compounds). After desalting and filtering to molecular weight *<* 250 Da, 300 representatives were selected by greedy max-min diversity on Morgan fingerprints, radius = 2, 2048 bits)^41^.

### 2.3 Boltz-2 co-folding with orthosteric blockers

Boltz-2 inputs were prepared in YAML format with fragment placement assessed under different paired co-folding conditions depending on task: a blocked arm in which an orthosteric pocket occupant was included as an additional co-folded entity, and an unblocked arm in which the fragment was co-folded with the protein alone. All other inputs were held constant between conditions, including the protein sequence, multiple sequence alignment (MSA), and fragment SMILES. Ten independent diffusion samples were generated per fragment per condition.

For kinase targets, the blocker was adenosine diphosphate (ADP), which occupies the conserved ATP-binding site and physically excludes test fragments from that volume during inference. ADP was chosen over ATP as the orthosteric placeholder because the absence of the *γ*-phosphate yielded more consistent convergence of the nucleotide pose across diffusion samples, providing more reproducible occlusion of the ATP-binding site. This is consistent with the higher representation of ADP relative to ATP in kinase crystal structures deposited in the Protein Data Bank (PDB)^38^, which likely reflects a stronger structural prior for the ADP-bound configuration in Boltz-2.

For the non-kinase targets, PTP1B and KRAS, the DADEpYL hexapeptide and GDP, respectively, served as the blocking chain in place of ADP (Section 2.1). All other protocol details were identical to the kinase screen.

### 2.4 Localization metrics

Fragment placement was quantified by heavy-atom center-of-mass (COM) distances in a common structural reference frame. Prior to the distance measurement, each predicted structure was aligned to the case-specific reference protein by C*α*-only superposition in PyMOL^42^, placing all predictions in the same coordinate frame as the orthosteric and allosteric reference ligands. Metrics were computed independently for each of the ten diffusion samples per fragment per condition and summarized as means across the ensemble. For allosteric localization, the distance was defined as the minimum Euclidean distance from the predicted fragment COM to the nearest heavy atom of any allosteric reference ligand in the per-kinase reference set:

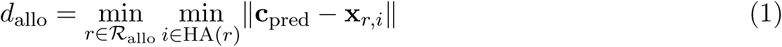

where *R*_allo_ is the set of all allosteric reference ligands for that kinase (the native parent-ligand coordinates of the construct plus all other panel allosteric structures for the same kinase), and HA(*r*) denotes the heavy atoms of reference ligand *r*. Orthosteric localization was measured analogously against the single canonical type-I reference ligand for each kinase (C*α*-superposed onto the parent crystal):

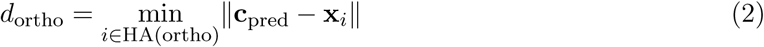

A fragment was classified as allosterically or orthosterically localized for a given sample if *d*_allo_ ≤ 5.0 Å or *d*_ortho_ ≤ 5.0 Å, respectively.

### 2.5 Absolute binding free energy calculations

Absolute binding free energy (ABFE) calculations were performed on 30 fragment–kinase complexes derived from ADP-blocked co-folding to assess whether predicted allosteric poses are thermodynamically viable. For two fragments (CDK-A017 and CDK-A026), calculations were additionally performed on orthosteric and cryptic poses to directly compare binding free energies across pocket environments for the same fragment. Input structures contained the kinase protein and the test fragment; the ADP blocker was removed prior to simulation. Protein coordinates were taken from C*α*-aligned Boltz outputs; fragment heavy-atom coordinates were preserved from the co-folded pose, with bond orders and explicit hydrogens assigned using the library SMILES as an RDKit^39^ template.

Proteins were parameterized with AMBER ff14SB^43^ and solvated in TIP3P explicit solvent^44^ at 0.15 M ionic strength. Ligands were parameterized with GAFF2^45^ and AM1-BCC partial charges. Hydrogen mass repartitioning (3.024 amu) enabled a 4 fs production timestep. Standard double-decoupling ABFE was performed with three thermodynamic legs: complex, solvent, and restraint. Boresch orientational restraints were applied in the bound state with harmonic force constants of 10 kcal mol^−1^ Å^−2^. Ligand decoupling used 16 equally spaced *λ* windows. Each leg followed energy minimization, NVT heating, and NPT equilibration prior to production. Production comprised 5.0 ns per complex and solvent leg and 2.5 ns for the restraint leg, run on GPU. The standard binding free energy was assembled as:

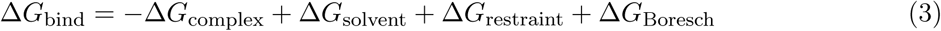

where Δ*G*_Boresch_ is the analytic correction for releasing orientational restraints in the bound state^34^. Free energies were estimated by MBAR^35^ at 298.15 K.

## 3 Results

### 3.1 CAFE universally restores non-orthosteric pocket exploration

To evaluate whether active-site occlusion provides a general, target-agnostic remedy for the prevalent orthosteric memorization using co-folding models, we established a multi-family benchmark comprising seven diverse targets: five representative human kinases (AKT2, CDK2, CHEK1, CSNK2A1, MAPK14), a protein tyrosine phosphatase (PTP1B), and a small GTPase (KRAS). Together, these systems span distinct fold topologies, catalytic mechanisms, and degrees of structural representation in the Protein Data Bank (PDB), ranging from less characterized pockets to heavily benchmarked targets like CDK2. We first quantify the non-orthosteric exploration rate of fragments defined as the percentage of diffusion samples in which the predicted fragment center-of-mass (COM) is at least 5.0 Å away from the canonical orthosteric pocket.

As shown in Table 1, active-site blocking increased non-orthosteric exploration of the native allosteric-derived fragments across the kinase and non-kinase panels. In the absence of blocking, most kinase targets showed strong fragment capture at the orthosteric site, with low non-orthosteric exploration rates for AKT2, CDK2, CHEK1, and MAPK14. The mean cross-kinase non-orthosteric exploration rate was 32.4%, but this average was notably elevated by CSNK2A1, which already showed high non-orthosteric exploration of fragments without ADP blocking; excluding CSNK2A1, the average no-blocker rate falls to 16.5%.

**Table 1:** Non-orthosteric exploration rate (%) under blocked and unblocked conditions. ADP is used as the blocker for all kinases. *n* denotes the number of allosteric-derived BRICS fragments evaluated per target.

| Target | Family | $n$ | No blocker | With blocker |
| --- | --- | --- | --- | --- |
| AKT2 | Human kinase | 15 | 6.0% | 100.0% |
| CDK2 | Human kinase | 38 | 8.2% | 99.7% |
| CHEK1 | Human kinase | 19 | 27.4% | 100.0% |
| CSNK2A1 | Human kinase | 11 | 96.4% | 100.0% |
| MAPK14 | Human kinase | 33 | 24.2% | 69.7% |
| PTP1B | Protein tyrosine phosphatase | 23 | 55.7% | 99.6% |
| KRAS | Small GTPase | 14 | 99.3% | 100.0% |

Physical blocking markedly disrupted this default preference for orthosteric binding across other target classes as well. For PTP1B, introducing a phosphotyrosine-containing peptide blocker elevated non-orthosteric exploration from 50.4% to 96.2%, demonstrating suppression comparable to the kinase family. In contrast, KRAS exhibited high non-orthosteric exploration even in the absence of a blocker (99.3% unblocked vs. 100.0% with GDP), consistent with a nucleotide-binding pocket that lacks the extreme structural overrepresentation driving memorization in kinases. These results indicate that when PDB training bias is minimal, such as for CSNK2A1 and KRAS, co-folding models naturally exercise good chemical reasoning regarding pocket exploration with fragments; CAFE recovers this latent capability in targets where heavy structural representation suppresses it. Supplementary Figure S2 reports the full distribution of fragment-to-orthosteric-pocket distances, with and without blocking, for all seven targets from Table 1.

We next broaden this analysis by co-folding the complete kinase-derived fragment library, all 116 orthosteric-derived and 116 allosteric-derived fragments, regardless of native origin, against each of the five kinases individually. Introducing ADP as a physical blocker consistently redirected both fragment cohorts across all kinases, elevating non-orthosteric exploration from 17.6% to 89.6% for allosteric-derived fragments, and from 5.7% to 80.4% for orthosteric-derived fragments (Figure 2A). Target-level analysis revealed that the magnitude of rescue reflects underlying PDB coverage while remaining universally effective. Unblocked CDK2 exhibited the lowest non-orthosteric exploration rate (2.2%), consistent with its standing as the most structurally characterized kinase in the PDB and thus its associated risk of training-data memorization. Despite this heavy bias, ADP blocking still drove a massive increase in non-orthosteric sampling (80.2%), confirming that physical occlusion of the canonical active site overcomes extreme training priors even if a minor residual orthosteric preference persists. Conversely, MAPK14 displayed a more moderate response to ADP blocking (52.5% non-orthosteric vs. the 85.0% cross-kinase mean), pointing to target-specific active-site pocket geometries that can partially accommodate co-folded fragments alongside the ADP blocker.

**Figure 2:**
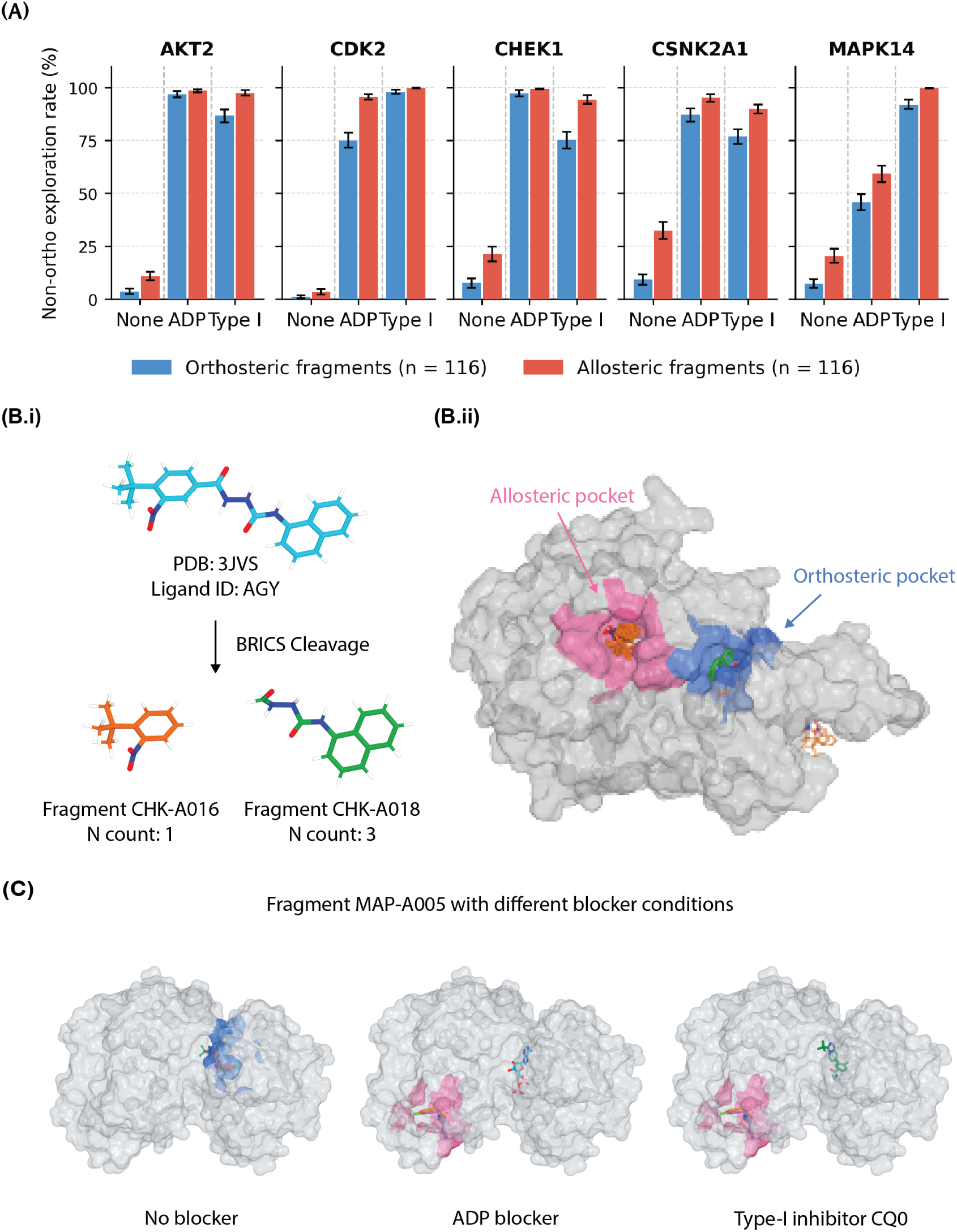
Active site blocking increases non-orthosteric exploration for both allosteric and orthosteric fragments across five kinases. (**A**) Non-orthosteric exploration rate (percentage of Boltz-2 samples with center-of-mass *>*5 Å from the orthosteric reference ligand center-of-mass) for orthosteric (blue, *n* = 116) and allosteric (red, *n* = 116) fragments under different blocking conditions: None, ADP, and Type I inhibitor. All fragments are co-folded against each kinase target regardless of native origin. Error bars represent standard error of the mean. **(B.i)** BRICS cleavage of the CHEK1 allosteric parent ligand AGY (PDB 3JVS) into fragments CHK-A016 and CHK-A018. **(B.ii)** Structural context of the orthosteric (blue) and allosteric (pink) pockets for this target. **(C)** Fragment MAP-A005 cofolded with no blocker, ADP blocker, and the Type-I inhibitor blocker CQ0. Using a blocker routes the fragment samples from the orthosteric site (blue) to the allosteric site (pink).

We find that fragment chemistry strongly predicted canonical ADP active-site capture across all five kinases. Specifically, aromatic ring count, hydrogen-bond acceptor count, and nitrogen atom count positively correlated with orthosteric placement (Spearman *ρ* ≥ 0.330, *p <* 0.0001; Supplementary Table S2, Figure S3). This reflects the the tridentate nitrogen pharmacophore of the conserved adenine moiety of ATP that routinely engage backbone hydrogen-bonding networks with kinase hinge residues (Supplementary Figure S4). Fragments capable of recapitulating this pharmacophore are pulled toward the orthosteric site regardless of their parent ligand’s binding mode. Fragments containing *N >* 2 exhibited significantly lower unblocked non-orthosteric exploration than those with *N* ≤ 2 (6.2% vs. 21.9%), demonstrating that orthosteric memorization is partially encoded in the fragment chemistry.

Figure 2B shows an example of this ligand-level memorization, in which Fragment CHK-A016 and Fragment CHK-A018 show drastically different pocket routing behaviors despite originating from the same allosteric parent ligand. All ten samples of the more nitrogen-rich CHK-A018 localize to the orthosteric pocket, while the majority of CHK-A016’s samples correctly localize to the allosteric pocket. Introducing ADP blocking effectively decoupled fragment chemistry from active-site capture, raising the cross-kinase mean non-orthosteric exploration rate to 85.0% across all fragment types and leaving hydrogen-bond donor count as the only weak residual correlation (Supplementary Table S2).

As evidence of the generality of CAFE, we find that using co-crystallized Type-I small-molecule inhibitors as orthosteric blockers also yielded excellent redirection away from the canonical active site, even raising the cross-kinase mean non-orthosteric exploration rate of all fragment types to 91.0% (Figure 2A and Supplementary Table S3). As illustrated in Figure 2C using fragment MAP-A005 co-folded with the ADP blocker versus the Type-I inhibitor blocker CQ0, confirms that synthetic co-crystal inhibitors can equivalently occlude active-site volume during joint diffusion beyond just natural cofactors such as ADP. Thus, while fragment chemistry contributes to orthosteric trapping, physical active-site occlusion serves as an essential, target-agnostic intervention that can override chemical bias and restore non-orthosteric pocket exploration, and is largely agnostic to the blocker used to do so.

### 3.2 CAFE targets canonical allosteric sites with high pose fidelity

Having established that physical active-site blocking forces fragments to explore outside of canonical orthosteric pockets, we next investigate whether each kinase’s native allosteric-derived fragments (the same population as in Table 1) selectively populate the functional allosteric sites of their parent ligands when blocked by ADP, using the spatial criteria defined previously. As shown in Figure 3A, on-target allosteric localization without the blocker was minimal across most kinases, with the exception of CSNK2A1 whose canonical allosteric pocket is intrinsically accessible without physical blocking. Introducing ADP blocking via CAFE produces marked increases in allosteric site occupancy for AKT2 and CSNK2A1, demonstrating that redirected fragments do not merely scatter randomly across the protein surface, but systematically funnel into established allosteric pockets. MAPK14 and CHEK1 exhibited a more heterogeneous response, with moderate increase in localization. Strikingly, CDK2 yielded near-zero canonical allosteric localization under both conditions despite achieving near-complete escape from the orthosteric site. This pronounced disconnect indicates that target-specific allosteric fragments redirected from the ATP site in CDK2 preferentially populate unannotated surface regions or cryptic pockets rather than canonical allosteric sites as illustrated in Supplementary Figure S2. As a complementary specific control, each kinase’s own orthosteric-derived fragments are also redirected into the allosteric pocket under blocking, though at lower rates on average than allosteric-derived fragments (Supplementary Figure S5).

**Figure 3:**
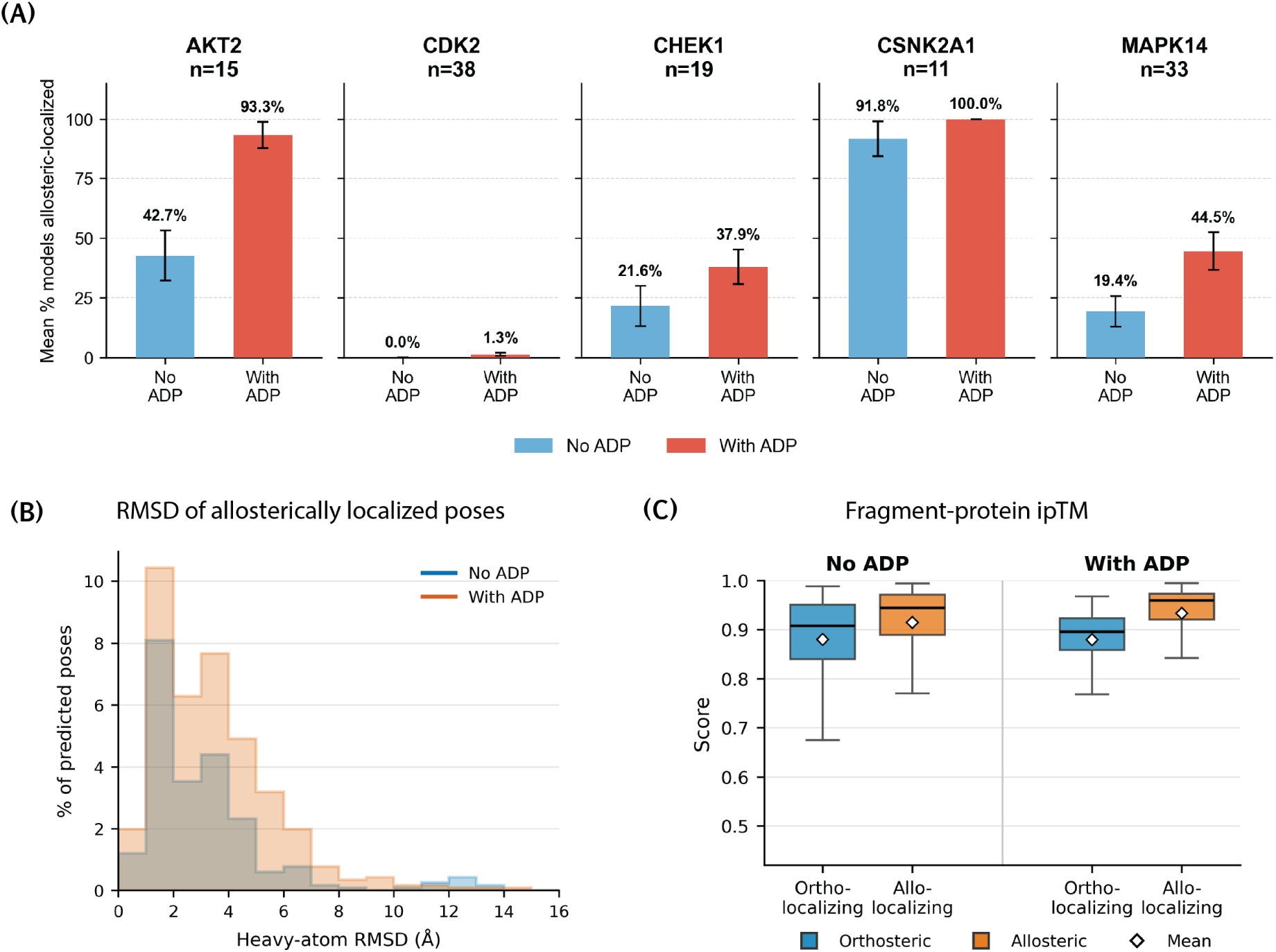
ADP blocking redirects individual fragments toward allosteric localization across five kinases. (**A**) Average percentage of native allosteric fragments with predicted center-of-mass within 5.0 Å of the nearest heavy atom of any allosteric reference ligand, without (blue) and with (red) ADP co-folding blocker. **(B)** Heavy-atom RMSD distribution of allosterically localizing poses relative to the crystallographic reference, with and without ADP blocking. **(C)** Fragment–protein ipTM scores for ortho-localizing versus allo-localizing poses, with and without ADP.

To corroborate the findings of localization rate with geometric data, we further show in Figure 3B the RMSD value between sampled fragment poses and their crystallographic-derived ones. In the distribution, those classified as localized show minimal RMSD changes indicating that samples are exploring within the allosteric pockets for different poses. On the other hand, other samples that did not land in the correct pocket shows more of a global surface exploration as the RMSD values reach 15.4 Å. We further evaluated whether Boltz-2’s internal fragment-protein ipTM confidence metric could serve as a reliable proxy for correct allosteric placement without explicit distance scoring (Figure 3C). In unblocked runs, allosterically localized poses were assigned negligibly higher fragment-protein ipTM than orthosterically localized ones (median 0.945 vs. 0.907). We note that ADP blocking did not significantly improve the Boltz-2 the ipTM confidence score between allosteric and orthosteric placements (0.959 vs. 0.896). In fact, the orthosteric and allosteric score distributions overlap almost entirely in both the blocked and unblocked conditions, making a high ipTM value for an individual pose uninformative about which pocket it occupies. This problem is also maintained with Boltz-2’s internal global confidence metric (Supplementary Table S5). Thus, native co-folding confidence scores cannot substitute for explicit geometric localization scoring in allosteric discovery screens.

### 3.3 Allosteric fragment poses show favorable binding free energies

To determine whether CAFE yields thermodynamically viable binding poses, we performed absolute binding free energy (ABFE) calculations across 30 fragment–kinase complexes spanning all five target kinases. Selected fragments represented cases exhibiting orthosteric localization in ≥ 60% of models in the absence of ADP but recovered the canonical allosteric pocket in at least one model upon blocking. This set was supplemented by fragments from AKT2 and CSNK2A1 that already engaged allosteric sites without ADP present, to broaden per-kinase coverage. For each fragment, Boltz-2 poses were used without post-processing or structural relaxation prior to the ABFE simulation, with only addition of hydrogens and charge assignment applied, and thus reflects the quality of the raw co-folding output directly. As shown in Figure 4 and Supplementary Table S4, the Boltz-predicted allosteric poses are energetically equivalent to or more favorable than the crystallographic reference for 21 of the 30 fragment–kinase complexes. Crucially, a Wilcoxon signed-rank test revealed no statistically significant difference between the overall binding free energy distributions (*p* = 0.22), confirming that CAFE-generated allosteric poses routinely achieve a thermodynamic quality comparable to experimental co-crystal structures.

**Figure 4:**
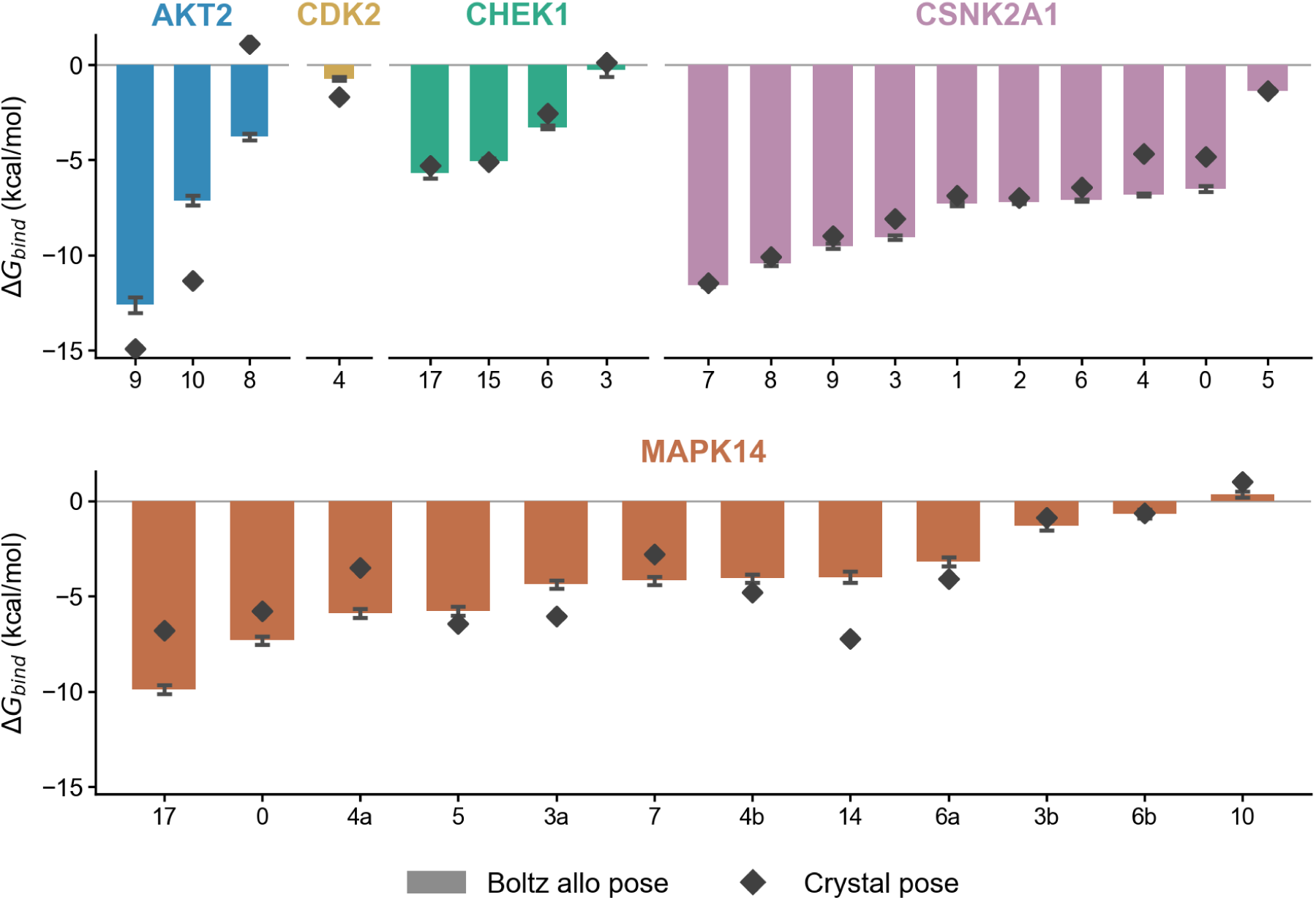
Boltz-predicted allosteric poses exhibit binding free energies comparable to experimental crystal poses (n=30). Bars show the Boltz allosteric-pose Δ*G_bind_*; diamond show the corresponding crystal-pose Δ*G_bind_*. Fragments are grouped by kinase and ordered by decreasing allosteric-pose favorability within each group. Fragments are labeled by the numeric suffix in Supplementary Table S1. When the same fragment was tested against two source PDB structures within a kinase (MAPK14 fragments 3, 4, and 6), “a”/“b” distinguish the two instances (a: 5N63, b: 5N64).

Behind this global equivalence, however, the thermodynamic agreement vary systematically based on target-specific pocket features and structural representation across the kinase panel. CSNK2A1 exhibited the strongest agreement, with all 10 fragment poses proving thermodynamically more favorable than their reference crystals (ΔΔ*G* ≤ 0), while CHEK1 similarly showed favorable binding energetics for 3 of 4 evaluated fragments. For MAPK14—the largest subset (*n* = 12) engaging a unique C^′^-lobe lipid-binding pocket distinct from the ATP site—we observed a balanced distribution of 7 allosteric-favorable and 5 crystal-favorable poses. Notably, this thermodynamic preference tracked strictly with parental crystal origin rather than fragment chemistry: all four fragments derived from parent PDB 5N63 favored the crystal reference, whereas all four derived from parent PDB 5N64 favored the Boltz pose.

This observation indicates that the observed variance reflects reference-specific crystallographic features, such as distinct loop conformations or water-mediated networks, rather than a systemic limitation of the co-folded pose. In contrast, targets with more limited representation displayed higher variability; AKT2 (*n* = 3) exhibited broad variance, producing both the single most allosteric-favorable pose in the panel and two of the least favorable. This divergence arises because the sole available AKT2 crystal reference (PDB 9C1W) contains zinc-mediated coordination contacts that standard non-covalent, restraint-based ABFE protocols are not designed to capture. Finally, for CDK2 (*n* = 1), candidate fragments routed predominantly to cryptic pockets rather than the canonical allosteric site under ADP blocking, precluding direct crystallographic comparison.

As a complementary validation, we evaluate the relative energetics of predicted allosteric versus orthosteric poses for fragments that were completely trapped in the orthosteric site (10*/*10 diffusion samples orthosteric without ADP; zero baseline allosteric occupancy) but gained allosteric engagement under ADP blocking. Allosteric poses were energetically equal to or more favorable than orthosteric poses for 5 of the 10 evaluated fragments, with no significant difference across the ensemble (Wilcoxon signed-rank test, *p* = 0.92). This demonstrates that physical redirection forces fragments into allosteric pockets without imposing energetic penalties relative to the active site. Furthermore, because each BRICS fragment represents a sub-component of a larger parent molecule, these results confirm that individual chemical moieties isolated by fragmentation harbor sufficient intrinsic affinity to drive site-specific allosteric recognition independently.

Importantly, this capacity for non-orthosteric redirection does not degrade the model’s fundamental chemical reasoning when active-site binding is the desired outcome. As a control, we applied the unblocked pipeline to INX-315, a clinical CDK2-selective inhibitor demonstrating greater than 200-fold biochemical preference over CDK6. The BRICS-derived heterocyclic core fragment successfully recapitulated this strict CDK2 selectivity via ABFE in the phosphorylated activation state, whereas two smaller peripheral fragments showed negligible binding to either target in isolation (Supplementary Table S5). This highlights that the CAFE framework natively preserves accurate orthosteric molecular recognition when it is the biologically relevant objective, but can instantly be flipped to unlock hidden allosteric real estate.

Additionally, the ability of the heterocyclic core to recapitulate target selectivity in the phosphorylated activation state, while the two peripheral fragments showed no meaningful binding independently, illustrates the anchor-and-elaborate logic central to FBDD. Fragments that contribute to selectivity in the context of a full drug may be invisible in isolation: peripheral contacts require a core anchor to position them productively. Identifying the selectivity-bearing anchor first and elaborating toward peripheral contacts second is the operationally correct order. The finding that unphosphorylated structures reversed the selectivity direction also carries a practical warning: activation state matching is not optional for fragment-level selectivity calculations, where every interaction is load-bearing. This sensitivity to conformational state is consistent with recent benchmarking showing that co-folding models exhibit severe mode collapse when predicting ligand-induced kinase conformational changes^22^.

### 3.4 CAFE uncovers novel cryptic pockets validated by fragment probes

Beyond recovering established allosteric sites, a frontier in structure-based drug discovery is the prospective identification of cryptic binding pockets, including transient surface cavities that are hard to sample in apo crystal structures but become more available in the holo induced-fit conformations. Traditional computational approaches for pocket identification are starkly polarized, relying either on rigid, geometry-based heuristics that miss transient states, or on all-atom molecular dynamics (MD) simulations that require a long time of intensive sampling^3^. By leveraging physical active-site blocking, CAFE provides an alternative by effectively re-purposing co-folding models into a pocket discovery tool guided by fragment-based probes that implicitly accounts for local receptor flexibility and induced-fit phenomena. Crucially, because these newly exposed pockets are mapped directly by the co-folded fragments, the fragments themselves could serve as potential fragment hits when paired with ABFE evaluations, allowing for simultaneous pocket and binder discovery.

To evaluate CAFE’s pocket discovery capability, we systematically analyze the population of sampled fragment poses that migrate into regions distinct from both the orthosteric site and canonical allosteric pockets. Across the five kinases, we identify 14 recurrent non-orthosteric, non-allosteric clusters using density-based spatial clustering (DBSCAN on fragment center-of-mass, *ɛ* = 5 Å, min_samples = 5). To benchmark these sites against conventional methods, we cross-reference them with pockets independently detected on static receptor structures using P2Rank^36^ and FPocket^37^. Overall, 10 of the 14 CAFE-uncovered clusters (71%) correspond to a cavity identified by either tools, but this recovery was highly non-uniform across the targets. While the CAFE-sampled pockets of CHEK1 (3/3), MAPK14 (2/2), and AKT2 (1/1) are fully captured by the static tools, only 4 of the 8 cryptic sites sampled by CDK2 fragments are detected by either P2Rank or FPocket. Furthermore, CAFE’s exclusive discovery of the two CDK2 pockets heavily populated by fragment probes underscores its prospective power, demonstrating that generative co-folding can successfully map highly plastic, non-orthosteric landscapes that remain inherently elusive to static, geometry-based prediction.

As shown in Figure 5A, the behavior of fragment CDK-A017 (PDB 8VQ3) demonstrates the unique pocket discovery capability of CAFE. Under ADP blocking, this fragment is successfully redirected to a cryptic pocket recovered by P2Rank but completely missed by FPocket. ABFE calculations further confirms that this co-folded cryptic pose is the most thermodynamically viable of all evaluated configurations (Δ*G* = −3.42 kcal/mol), significantly outperforming both its sampled orthosteric pose (+1.42 kcal/mol) and the experimental crystal-native reference pose (−0.36 kcal/mol). A second case, fragment CDK-A026 (PDB 8VQ4), demonstrates the utility of this approach when a classical crystal-pose comparison is fundamentally impossible. As a peripheral BRICS fragment, its crystallographically derived pose retains virtually none of the parent molecule’s core protein contacts. Despite that, CAFE successfully routes the fragment to a cryptic orientation that is more favorable (Δ*G* = −3.83 kcal/mol) than the sampled orthosteric alternative (−2.97 kcal/mol), further proving the viability of this newly exposed pocket. Together, these cases show that ADP-blocked co-folding has the potential to drive fragments into alternative pockets with overall desirable stability proven by free energy.

**Figure 5:**
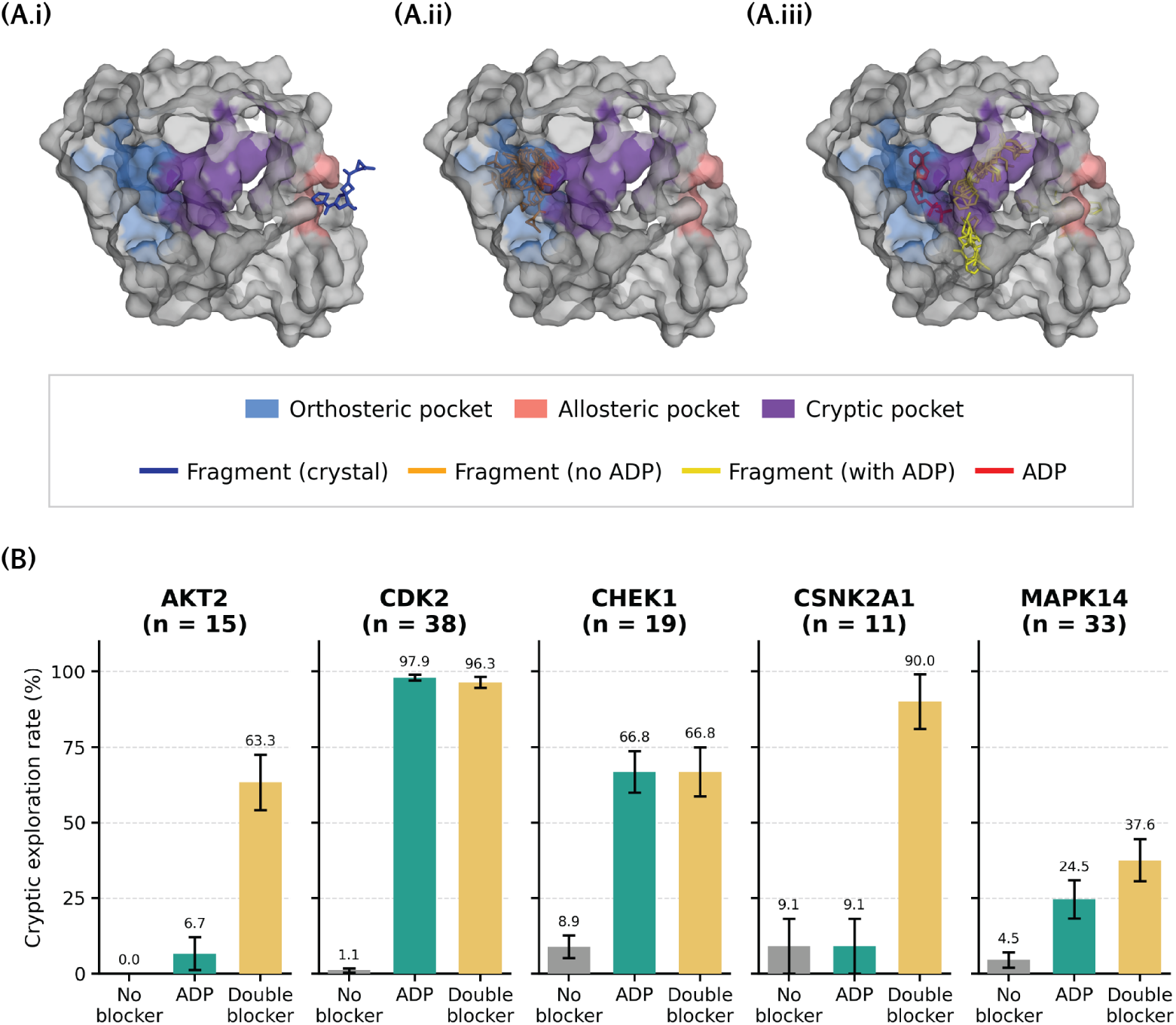
ADP blocking redirects a CDK2 fragment to a cryptic pocket undetected by conventional pocket-prediction tools, and pairing it with an allosteric blocker further enriches cryptic-site sampling. CDK-A017 (PDB: 8VQ3). **(A.i)** Reference ligands and predicted pocket sur-faces. Cryptic (purple) pocket recovered by P2Rank but not FPocket. **(A.ii)** Unblocked pose, sampling the orthosteric pocket (Δ*G* = +1.42 kcal/mol). **(A.iii)** ADP-blocked pose, redirected to the cryptic pocket and thermodynamically favored over both the orthosteric (Δ*G* = +1.42 kcal/mol) and crystal-native (Δ*G* = −0.36 kcal/mol) poses (Δ*G* = −3.42 kcal/mol). **(B)** Cryptic exploration rate (samples localized to neither orthosteric nor allosteric sites) across five kinases under no blocker, ADP alone, and ADP plus the parent allosteric ligand (double blocker). Error bars: SEM. The double blocker raises cryptic exploration beyond ADP alone for AKT2, CSNK2A1, and MAPK14, but not CDK2 or CHEK1, where cryptic sampling is already saturated under ADP alone.

Because ADP blocking alone can still permit fragments to populate the canonical allosteric site rather than cryptic pockets specifically, we next tested whether jointly occluding both the orthosteric pocket and the canonical allosteric pocket could deliberately steer fragments toward cryptic sites. In this double-blocker condition, ADP and the parent allosteric ligand were co-folded together as competing occupants, leaving only cryptic and other non-canonical surface regions available to the fragment. As shown in Figure 5B, this strategy increased cryptic pocket exploration substantially beyond ADP blocking alone for three of the five kinases: AKT2 (6.7% → 63.3%), CSNK2A1 (9.1% → 90.0%), and MAPK14 (24.5% → 37.6%). CDK2, which already showed near-saturating cryptic exploration under ADP blocking alone (97.9%), was unaffected by the additional allosteric block (96.3%), consistent with its canonical allosteric pocket already being functionally non-competitive for fragment occupancy in this system. CHEK1 likewise showed no further shift (66.8% under both conditions), indicating that occlusion of the orthosteric pocket alone was already sufficient to saturate cryptic-site sampling for this target. Together, these results show that CAFE’s site-selection behavior can be tuned further at inference time: pairing the orthosteric blocker with a canonical allosteric occupant provides a targeted mechanism for enriching cryptic pocket discovery specifically in kinases where the annotated allosteric site would otherwise compete with cryptic pockets for fragment occupancy, while adding no benefit where that competition is already minimal.

### 3.5 CAFE as a screening strategy using commercial fragment libraries

Fragment-based drug discovery (FBDD) offers an exceptionally efficient route to scan chemical space for challenging targets, but its real-world success hinges on identifying cryptic or allosteric binding modes early enough to guide whole-ligand expansion^31^. To bridge this gap, we present a prospective screening recipe using CAFE as a rapid, dynamic launchpad to discover, score, and prioritize non-orthosteric hits from unannotated commercial libraries. To demonstrate this workflow in a prospective setting, we screened 300 chemically diverse fragments selected from the Enamine High Fidelity Fragment Library^46^ against each of our five target kinases under ADP blocking conditions.

As demonstrated in Figure 6, this prospective workflow operates as a tiered funnel consisting of three major stages: screening, down-selection, and validation. First, CAFE samples candidate fragment poses and potential cryptic pockets across the library screen. Following this initial phase, a canonical allosteric pocket or a newly identified cryptic site is designated as the pocket of interest, against which the generated poses are systematically scored based on their protein–ligand interaction counts via the Protein-Ligand Interaction Profiler (PLIP)^47^. Based on these rankings, the top 15 candidate fragments per pocket are down-selected for explicit absolute binding free energy (ABFE) evaluation. During this validation stage, any poses in which the fragment drifts out of its intended pocket over the course of the 5-ns simulations are excluded. The remaining structurally stable poses with favorable thermodynamic profiles are prioritized as validated fragment anchors for downstream whole-ligand design.

**Figure 6:**
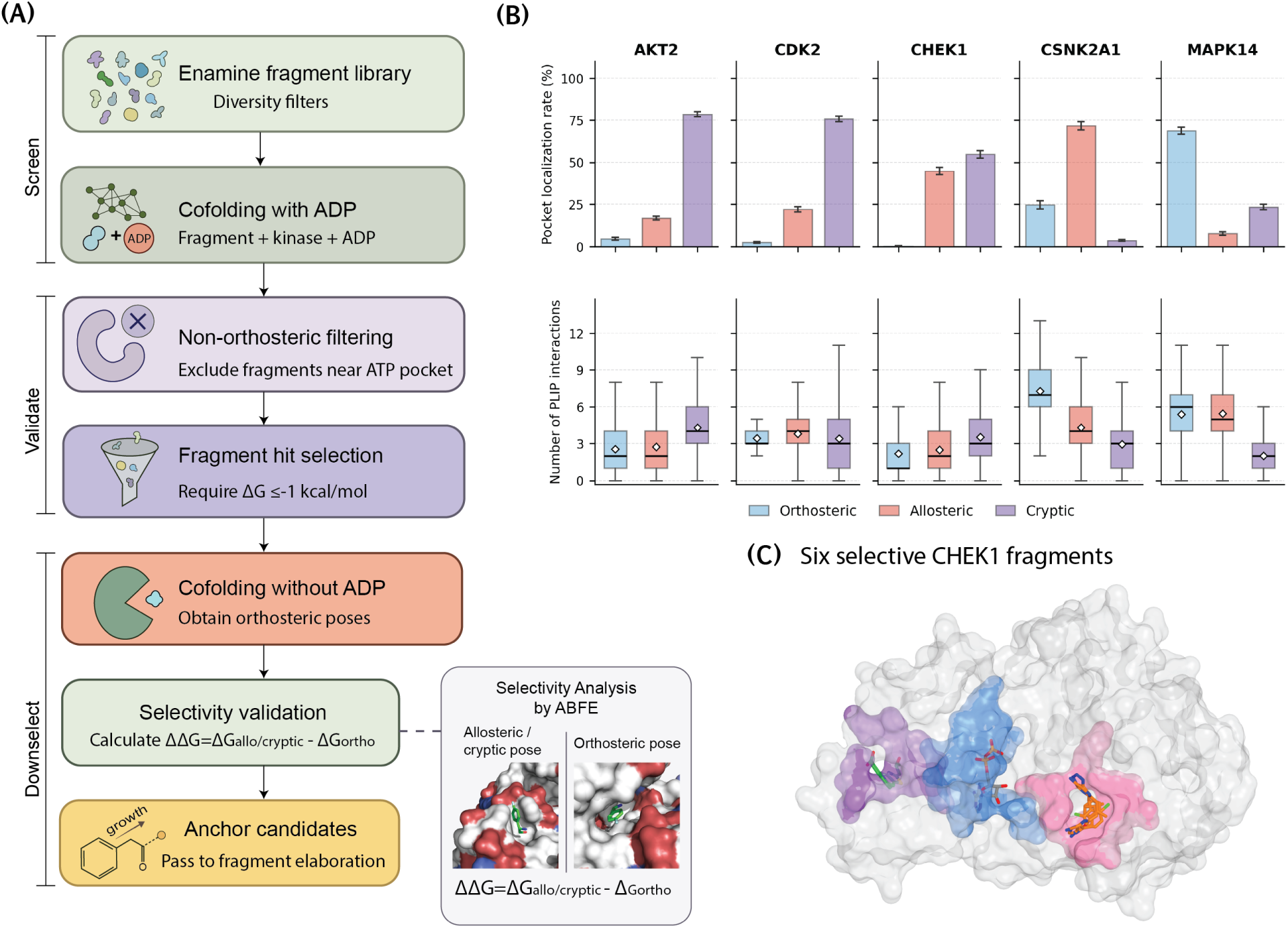
The CAFE prospective screening workflow and pocket-selective fragment recovery. (**A**) Screening funnel: Enamine fragments are co-folded with each kinase under ADP blocking, filtered for non-orthosteric localization, down-selected by Δ*G* ≤ −1 kcal/mol, then re-folded without ADP and validated by paired ABFE (ΔΔ*G* = Δ*G*_allo/cryptic_ −Δ*G*_ortho_) to confirm blocker-dependent selectivity. **(B)** Per-kinase pocket localization rate (top) and PLIP interaction counts (bottom) for fragments assigned to orthosteric, allosteric, and cryptic sites. **(C)** Structural example of six CHEK1 fragments selectively engaging allosteric (pink), and cryptic (purple) pockets, with the ADP blocker present in the orthosteric pocket (blue)

To evaluate the prospective utility of the CAFE platform, we initiate a screening campaign targeting the five kinase targets, focusing on one canonical allosteric site and one newly identified cryptic site per target. Following filtering and down-selection, 107 fragments are retained for the final ABFE evaluations. 33 of these fragments (31%) emerge as thermodynamically favorable hits (Δ*G* ≤ −1 kcal/mol) spanning all five kinases (AKT2: 4, CDK2: 5, CHEK1: 6, CSNK2A1: 7, MAPK14: 11) (Supplementary Table S6). This robust hit rate underscores CAFE’s capability as a prospective discovery engine to isolate viable starting matter directly from generic, independently sourced commercial collections. As a critical quality-control step, we re-evaluate the 33 favorable hits by generating independent diffusion trajectories in the complete absence of the active-site blocker. Strikingly, 30 of the 33 hits (91%) immediately revert to orthosteric localization; for 16 of these fragments, all ten independent trajectories route exclusively into the orthosteric pocket. This clean reversal demonstrates that the observed allosteric and cryptic engagement is strictly attributable to the tunable ADP-blocking mechanism, rather than a systemic bias in fragment selection or pose scoring.

To prioritize these validated hits for downstream whole-ligand expansion, we next evaluate their energetic selectivity by executing parallel ABFE calculations on their corresponding orthosteric configurations (Supplementary Table S6). Among the 30 confirmed fragments, 23 yield orthosteric poses that remain structurally stable throughout the simulation, enabling a direct, paired thermodynamic comparison. Crucially, the non-orthosteric configurations are energetically equivalent to or more favorable than their competitive orthosteric alternatives for 14 of the 23 fragments (61%), out-performing them with a substantial mean Δ*G* gap of −2.10 vs. −0.62 kcal/mol. Collectively, these findings transition CAFE from a validation concept into an actionable prospective asset: by simultaneously generating structural coordinates and direct thermodynamic validation, CAFE provides an efficient, accessible launchpad for translating fragment libraries into selective, non-orthosteric chemical leads.

## 4 Discussion and Conclusion

Fragment-based drug discovery occupies a privileged position in early-stage hit identification: fragments sample chemical space more efficiently than drug-like molecules, bind with higher ligand efficiency, and reveal pharmacophoric hot spots that can be elaborated into leads with superior selectivity profiles. However, the utility of FBDD for allosteric target engagement remains limited by the difficulty of directing fragments away from dominant orthosteric sites. The present work addresses this limitation by introducing orthosteric-blocked co-folding as a structurally grounded, computationally lightweight strategy for redirecting fragment exploration toward the non-orthosteric pocket landscape. By treating the orthosteric occupant as an additional co-folded entity, the method physically excludes fragments from the canonical binding site during inference, leveraging the joint diffusion process that makes co-folding models well suited to capturing induced-fit effects at the fragment level. The approach requires no experimental pocket detection, no docking grid preparation, and no post-hoc structural refinement, yet recovers allosteric and cryptic binding geometries of thermodynamic quality comparable to crystallographic references. Applied across kinases spanning a wide range of PDB coverage using both ADP and type I ligands as orthosteric blockers, and extension to non-kinase targets, the CAFE method demonstrates consistent suppression of orthosteric memorization and reliable redirection of fragment sampling to non-canonical sites.

Using Boltz-2 as the co-folding engine, orthosterically-localized samples consistently received higher confidence scores than allosterically-localized ones, and this bias persisted under ADP blocking at the fragment–protein interface level. Practitioners using confidence-ranked pose selection without explicit localization scoring would systematically prefer orthosteric placements even when the majority of poses have been successfully redirected. Explicit center-of-mass localization scoring against crystallographic reference pockets is therefore a necessary complement to internal model confidence metrics in allosteric fragment screening workflows. This finding echoes the broader observation that co-folding models largely recapitulate training data geometry^16,17^, and that non-trivial pose quality requires structural priors that go beyond model confidence alone.

But we also emphasize that the Boltz-2 engine was also capable of excellent chemical reasoning when CAFE diverted fragments to allosteric or cryptic binding sites, indicating inherent capabilities that were unleashed under steric blocking. A central finding is that ABFE calculations on raw Boltz-2-predicted poses without structural relaxation, refinement, or pose optimization return binding free energies comparable to or exceeding those from crystallographically derived reference poses. This is a non-trivial result: it implies that the joint protein–ligand diffusion process in Boltz-2 produces physically reasonable binding geometries even for fragments, where the signal-to-noise ratio for pose determination is inherently low. In several cases, co-folded poses were substantially more favorable than crystal references, suggesting that joint diffusion can sample binding geometries not captured in parent co-crystal structures. The absence of a refinement step also means that co-folding output can be fed directly into an ABFE pipeline, collapsing what is typically a multi-stage structure preparation workflow into a single inference step.

The cryptic pocket results represent the most drug-discovery-relevant outcome of the CAFE framework. While non-orthosteric, non-allosteric clusters identified under ADP blocking for the kinases can often corresponded to a pocket independently detected by either P2Rank or FPocket, CAFE found additional new cryptic sites that are thermodynamically favorable. A logical extension of CAFE was to introduce both ADP and allosteric blockers to force the discovery of new cryptic binding sites that also showed fragments with favorable thermodynamic binding. Prospective screening of an independent, chemically diverse commercial fragment library extended this result across all five kinases in which a third of screened fragments were thermodynamically favorable under ADP blocking compared to the orthosteric site. Where a paired orthosteric comparison was possible, the majority of non-orthosteric poses yielded ABFEs that were equal to or more favorable, confirming that the cryptic and allosteric pockets identified by ADP-blocked co-folding are not artifacts of the curated validation library.

Looking forward, the natural next step is to connect fragment hits to an elaboration pipeline. Tools such as Fragmenstein^48^ for fragment merging and growing, and LinkLlama^49^ for fragment linking, provide complementary routes to converting fragment anchors into lead-like molecules while preserving the binding geometry identified by blocked co-folding. Re-co-folding of elaborated candidates with ADP blocking provides a rapid computational check that the non-orthosteric binding mode is retained upon growth. For cryptic pockets in particular, where pocket geometry is not pre-organized and every interaction is load-bearing, agentic fragment design workflows in which a generative model iteratively proposes, evaluates, and refines candidates against explicit ABFE feed-back may be better suited than enumeration-based approaches to navigating the narrow tolerance for suboptimal contacts. The pocket descriptor framework developed here, connecting hydrophobic fraction, burial depth, and charged residue content to ABFE-predicted bindability, provides a natural scoring function for guiding such agentic loops toward pockets likely to reward elaboration.

Finally, although Boltz-2 was used as the co-folding engine, the blocking strategy is model-agnostic in principle. Any joint protein–ligand structure prediction model that accepts multi-chain inputs — including AlphaFold3, RoseTTAFold All-Atom, or future co-folding architectures — could implement the same orthosteric (or allosteric) exclusion logic by including an appropriate pocket occupant as an additional chain. For harder targets, where co-folding models struggle to assemble the poses of the protein based on sequences which lack sufficient structural representation in the PDB, CAFE will also struggle. But we expect co-folding models to improve in ligand pose accuracy and conformational sampling, such that the quality of redirected fragment placements using CAFE are expected to improve correspondingly. The CAFE framework thus provides the blueprint that can be adopted across the evolving landscape of structure prediction models, with the blocking strategy itself remaining unchanged regardless of which model is used.

## 5 Supplementary information

Details of the data formats and usage notes are provided in the Supplementary Information.

## 6 Data and code availability

The CAFE code and fragment library datasets used in this paper are available at https://github.com/THGLab/CAF

## 7 Author contributions

All authors conceptualized the scientific goal of the project. JP wrote the code. All authors performed analysis for the results section, and wrote and editted the manuscript.

## 9 Acknowledgments

We thank Yingze Wang for helpful discussions. This work was supported by National Institute of Allergy and Infectious Disease grant U19-AI171954. This research used computational resources of the National Energy Research Scientific Computing, a DOE Office of Science User Facility supported by the Office of Science of the U.S. Department of Energy under Contract No. DE-AC02-05CH11231.

