## Supplementary material for "CAFE: A Co-folding Approach for Fragment Exploration of Allosteric and Cryptic Binding Sites": SI

#### 1 Supplementary Tables

**Table S1:** Fragment library catalog consisting of 116 allosteric and 116 orthosteric fragments.

| Fragment ID | PDB | SMILES |
|---|---|---|
| AKT-A000 | 8Q61 | CC1(O)CC(N)(c2ccccc2)C1 |
| AKT-A001 | 8Q61 | CCCN1C(=O)COc2nc(-c3ccc(C4(N)CC(C)(O)C4)cc3)ccc21 |
| AKT-A002 | 8Q61 | CCCN1C(=O)COc2nc(-c3ccccc3)c(-c3ccccc3)cc21 |
| AKT-A003 | 8Q61 | CCCN1C(=O)COc2ncc(-c3ccccc3)cc21 |
| AKT-A004 | 9C1W | CCNCc1ccc(-n2c(-c3cccnc3N)nc3ccc(-c4ccccc4)nc32)cc1 |
| AKT-A005 | 9C1W | CCc1ccc(C=O)c(O)c1 |
| AKT-A006 | 9C1W | CNCCc1ccc(C=O)c(O)c1 |
| AKT-A007 | 9C1W | Cc1ccc(-n2c(-c3cccnc3N)nc3ccc(-c4ccccc4)nc32)cc1 |
| AKT-A008 | 9C1W | NCCc1ccc(C=O)c(O)c1 |
| AKT-A009 | 9C1W | NCc1ccc(-n2c(-c3cccnc3N)nc3ccc(-c4ccccc4)nc32)cc1 |
| AKT-A010 | 9C1W | Nc1ncccc1-c1nc2ccc(-c3ccccc3)nc2[nH]1 |
| AKT-A011 | 9C1W | Nc1ncccc1-c1nc2ccc(-c3ccccc3)nc2n1-c1ccccc1 |
| AKT-A012 | 9C1W | Nc1ncccc1-c1nc2cccnc2n1-c1ccc(CNCCc2ccc(C=O)c(O)c2)cc1 |
| AKT-A013 | 9C1W | O=Cc1ccc(CCNCc2ccc(-n3cnc4ccc(-c5ccccc5)nc43)cc2)cc1O |
| AKT-A014 | 9C1W | O=Cc1ccc(CCNCc2ccccc2)cc1O |
| AKT-O000 | 2JDO | CCNCCNS(=O)(=O)c1cccc2cnccc12 |
| AKT-O001 | 2JDO | CCNS(=O)(=O)c1cccc2cnccc12 |
| AKT-O002 | 2JDO | COCCNCCNS(=O)(=O)c1cccc2cnccc12 |
| AKT-O003 | 2JDO | NCCNCCOCc1ccc(Cl)cc1 |
| AKT-O004 | 2JDO | NCCNS(=O)(=O)c1cccc2cnccc12 |
| AKT-O005 | 2JDO | O=S(=O)(NCCNCCO)c1cccc2cnccc12 |
| AKT-O006 | 3D0E | CC(C)(O)C#Cc1ncc(OCC2CCCNC2)c2[nH]c(-c3nonc3N)nc12 |
| AKT-O007 | 3D0E | CCn1c(-c2nonc2N)nc2c(C#CC(C)(C)O)ncc(O)c21 |
| AKT-O008 | 3D0E | CCn1c(-c2nonc2N)nc2c(C#CC(C)(C)O)ncc(OC)c21 |
| AKT-O009 | 3D0E | CCn1c(-c2nonc2N)nc2c(C#CC(C)(C)O)nccc21 |
| AKT-O010 | 3D0E | CCn1cnc2c(C#CC(C)(C)O)ncc(OCC3CCCNC3)c21 |
| AKT-O011 | 3E88 | CC(C)(O)C#Cc1nc(OCC(N)Cc2ccccc2)cc2[nH]c(-c3nonc3N)nc12 |
| AKT-O012 | 3E88 | CCn1c(-c2nonc2N)nc2c(C#CC(C)(C)O)nc(O)cc21 |
| AKT-O013 | 3E88 | CCn1c(-c2nonc2N)nc2c(C#CC(C)(C)O)nc(OCC(C)N)cc21 |
| AKT-O014 | 3E88 | CCn1cnc2c(C#CC(C)(C)O)nc(OCC(N)Cc3ccccc3)cc21 |
| CDK-A000 | 6Q3F | Nc1cc(Br)ccn1 |
| CDK-A001 | 6Q49 | O=c1cc(Br)cc[nH]1 |
| CDK-A002 | 6Q4K | Cc1cccc(C(=O)Nc2ccc(Cl)cc2)c1 |
| CDK-A003 | 6Q4K | NC(=O)c1cccc(C=CC(=O)O)c1 |
| CDK-A004 | 6Q4K | O=CNc1ccc(Cl)cc1 |
| CDK-A005 | 6Q4K | O=Cc1cccc(C=CC(=O)O)c1 |
| CDK-A006 | 8VQ3 | CC(CC(=O)N1CCOCC1)OCC1CCCCC1 |
| CDK-A007 | 8VQ3 | CC(O)C(NC(=O)C1CN(C(=O)c2cncs2)CC12CN(C(=O)C1CC1(C)C)C2)C (=O)N1CCOCC1 |
| CDK-A008 | 8VQ3 | CC(OCC1CCCCC1)C(C=O)NC(=O)C1CN(C(=O)c2cncs2)CC12CN(C(=O)C 1CC1(C)C)C2 |

Table S1 – continued from previous page

| Fragment ID | PDB | SMILES |
|---|---|---|
| CDK-A009 | 8VQ3 | CC(OCC1CCCCC1)C(N)C(=O)N1CCOCC1 |
| CDK-A010 | 8VQ3 | CC(OCC1CCCCC1)C(NC(=O)C1CN(C(=O)c2cncs2)CC12CN(C=O)C2)C(=O)N1CCOCC1 |
| CDK-A011 | 8VQ3 | CC(OCC1CCCCC1)C(NC(=O)C1CN(C(=O)c2cncs2)CC12CNC2)C(=O)N1CCOCC1 |
| CDK-A012 | 8VQ3 | CC(OCC1CCCCC1)C(NC(=O)C1CN(C=O)CC12CN(C(=O)C1CC1(C)C)C2)C(=O)N1CCOCC1 |
| CDK-A013 | 8VQ3 | CC(OCC1CCCCC1)C(NC(=O)C1CNCC12CN(C(=O)C1CC1(C)C)C2)C(=O)N1CCOCC1 |
| CDK-A014 | 8VQ3 | CC(OCC1CCCCC1)C(NC=O)C(=O)N1CCOCC1 |
| CDK-A015 | 8VQ3 | CC1(C)CC1C(=O)N1CC2(CCN(C(=O)c3cncs3)C2)C1 |
| CDK-A016 | 8VQ3 | CC1(C)CC1C(=O)N1CC2(CN(C(=O)c3cncs3)CC2C(N)=O)C1 |
| CDK-A017 | 8VQ3 | CC1(C)CC1C(=O)N1CC2(CN(C(=O)c3cncs3)CC2C=O)C1 |
| CDK-A018 | 8VQ3 | CCC(NC(=O)C1CN(C(=O)c2cncs2)CC12CN(C(=O)C1CC1(C)C)C2)C(=O)N1CCOCC1 |
| CDK-A019 | 8VQ3 | COC(C)C(NC(=O)C1CN(C(=O)c2cncs2)CC12CN(C(=O)C1CC1(C)C)C2)C(=O)N1CCOCC1 |
| CDK-A020 | 8VQ4 | CC(OCCC1CCCCC1)C(C=O)NC(=O)C1CN(C(=O)c2cnn(Cc3ccccc3)c2)CC12CN(C(=O)C1CC1(C)C)C2 |
| CDK-A021 | 8VQ4 | CC1(C)CC1C(=O)N1CC2(CCN(C(=O)c3cnn(Cc4ccccc4)c3)C2)C1 |
| CDK-A022 | 8VQ4 | CC1(C)CC1C(=O)N1CC2(CN(C(=O)c3cnn(Cc4ccccc4)c3)CC2C(N)=O)C1 |
| CDK-A023 | 8VQ4 | CC1(C)CC1C(=O)N1CC2(CN(C(=O)c3cnn(Cc4ccccc4)c3)CC2C=O)C1 |
| CDK-A024 | 8VQ4 | CCC(NC(=O)C1CN(C(=O)c2cnn(Cc3ccccc3)c2)CC12CN(C(=O)C1CC1(C)C)C2)C(=O)NC |
| CDK-A025 | 8VQ4 | CCOC(C)C(NC(=O)C1CN(C(=O)c2cnn(Cc3ccccc3)c2)CC12CN(C(=O)C1CC1(C)C)C2)C(=O)NC |
| CDK-A026 | 8VQ4 | CNC(=O)C(N)C(C)OCCC1CCCCC1 |
| CDK-A027 | 8VQ4 | CNC(=O)C(NC(=O)C1CN(C(=O)c2cn[nH]c2)CC12CN(C(=O)C1CC1(C)C)C2)C(C)OCCC1CCCCC1 |
| CDK-A028 | 8VQ4 | CNC(=O)C(NC(=O)C1CN(C(=O)c2cnn(C)c2)CC12CN(C(=O)C1CC1(C)C)C2)C(C)OCCC1CCCCC1 |
| CDK-A029 | 8VQ4 | CNC(=O)C(NC(=O)C1CN(C(=O)c2cnn(Cc3ccccc3)c2)CC12CN(C(=O)C1CC1(C)C)C2)C(C)O |
| CDK-A030 | 8VQ4 | CNC(=O)C(NC(=O)C1CN(C(=O)c2cnn(Cc3ccccc3)c2)CC12CN(C=O)C2)C(C)OCCC1CCCCC1 |
| CDK-A031 | 8VQ4 | CNC(=O)C(NC(=O)C1CN(C(=O)c2cnn(Cc3ccccc3)c2)CC12CNC2)C(C)OCCC1CCCCC1 |
| CDK-A032 | 8VQ4 | CNC(=O)C(NC(=O)C1CN(C=O)CC12CN(C(=O)C1CC1(C)C)C2)C(C)OCCC1CCCCC1 |
| CDK-A033 | 8VQ4 | CNC(=O)C(NC(=O)C1CNCC12CN(C(=O)C1CC1(C)C)C2)C(C)OCCC1CCCCC1 |
| CDK-A034 | 8VQ4 | CNC(=O)C(NC=O)C(C)OCCC1CCCCC1 |
| CDK-A035 | 8VQ4 | CNC(=O)CC(C)OCCC1CCCCC1 |
| CDK-A036 | 8VQ4 | O=Cc1cnn(Cc2ccccc2)c1 |
| CDK-A037 | 8VQ4 | c1ccc(Cn2cccn2)cc1 |
| CDK-O000 | 3QTX | NS(=O)(=O)c1ccccc1 |
| CDK-O001 | 3QTX | Nc1ccc(S(N)(=O)=O)cc1 |
| CDK-O002 | 3QTX | Nc1csc(Nc2ccc(S(N)(=O)=O)cc2)n1 |
| CDK-O003 | 3QTX | Nc1nc(N)c(C(=O)c2cccc([N+](=O)[O-])c2)s1 |
| CDK-O004 | 3QTX | Nc1nc(Nc2ccc(S(N)(=O)=O)cc2)sc1C=O |
| CDK-O005 | 3QTX | Nc1ncsc1C(=O)c1cccc([N+](=O)[O-])c1 |
| CDK-O006 | 3QTX | O=Cc1cccc([N+](=O)[O-])c1 |
| CDK-O007 | 3RNI | Nc1cccc(S(N)(=O)=O)c1 |
| CDK-O008 | 3RNI | Nc1csc(Nc2cccc(S(N)(=O)=O)c2)n1 |
| CDK-O009 | 3RNI | Nc1nc(N)c(C(=O)c2ccccc2)s1 |
| CDK-O010 | 3RNI | Nc1nc(Nc2cccc(S(N)(=O)=O)c2)sc1C=O |
| CDK-O011 | 3RNI | Nc1ncsc1C(=O)c1ccccc1 |
| CDK-O012 | 4GCJ | Nc1nc(N)c(C(=O)c2ccccc2[N+](=O)[O-])s1 |
| CDK-O013 | 4GCJ | Nc1ncsc1C(=O)c1ccccc1[N+](=O)[O-] |
| CDK-O014 | 4GCJ | O=Cc1ccccc1[N+](=O)[O-] |
| CDK-O015 | 8ERD | CC1CCCN1C(=O)c1ccc2cnc(N)nc2c1 |

Table S1 – continued from previous page

| Fragment ID | PDB | SMILES |
|---|---|---|
| CDK-O016 | 8ERD | CC1CCCN1C(=O)c1ccc2cnc(NC3CCNCC3)nc2c1 |
| CDK-O017 | 8ERD | CC1CCCN1C(=O)c1ccc2cncnc2c1 |
| CDK-O018 | 8ERD | CS(=O)(=O)N1CCC(N)CC1 |
| CDK-O019 | 8ERD | CS(=O)(=O)N1CCC(Nc2ncc3ccc(C=O)cc3n2)CC1 |
| CDK-O020 | 8ERD | CS(=O)(=O)N1CCC(Nc2ncc3ccccc3n2)CC1 |
| CDK-O021 | 8ERD | CS(=O)(=O)N1CCCCC1 |
| CDK-O022 | 8ERN | CNS(=O)(=O)c1ccc(N)c(C)c1 |
| CDK-O023 | 8ERN | CNS(=O)(=O)c1ccc(Nc2ncc3ccc(C=O)cc3n2)c(C)c1 |
| CDK-O024 | 8ERN | CNS(=O)(=O)c1ccc(Nc2ncc3ccccc3n2)c(C)c1 |
| CDK-O025 | 8ERN | CNS(=O)(=O)c1cccc(C)c1 |
| CDK-O026 | 8ERN | Cc1cc([SH](=O)=O)ccc1Nc1ncc2ccc(C(=O)N3CCCC3C)cc2n1 |
| CHK-A000 | 3F9N | C1CNCC2(C1)CCCNC2 |
| CHK-A001 | 3F9N | CCCCC(C=O)Sc1nc2ccccc2c(=O)n1-c1cccc(Cl)c1 |
| CHK-A002 | 3F9N | CCCCC(S)C(=O)N1CCCC2(CCCNC2)C1 |
| CHK-A003 | 3F9N | CCCCC(Sc1nc2ccccc2c(=O)[nH]1)C(=O)N1CCCC2(CCCNC2)C1 |
| CHK-A004 | 3F9N | CCCCCC(=O)N1CCCC2(CCCNC2)C1 |
| CHK-A005 | 3F9N | O=c1c2ccccc2nc(S)n1-c1cccc(Cl)c1 |
| CHK-A006 | 3F9N | O=c1c2ccccc2ncn1-c1cccc(Cl)c1 |
| CHK-A007 | 3JVR | CC(O)c1nc2ccccc2[nH]1 |
| CHK-A008 | 3JVR | CC(OC(N)=O)c1nc2ccccc2[nH]1 |
| CHK-A009 | 3JVR | CC(OC=O)c1nc2ccccc2[nH]1 |
| CHK-A010 | 3JVR | CCOC(=O)Nc1ccc(Cl)c(Cl)c1 |
| CHK-A011 | 3JVR | Nc1ccc(Cl)c(Cl)c1 |
| CHK-A012 | 3JVR | O=C(O)Nc1ccc(Cl)c(Cl)c1 |
| CHK-A013 | 3JVR | O=CNc1ccc(Cl)c(Cl)c1 |
| CHK-A014 | 3JVS | CC(C)(C)c1ccc(C(=O)NNC(N)=O)cc1[N+](=O)[O-] |
| CHK-A015 | 3JVS | CC(C)(C)c1ccc(C(=O)NNC=O)cc1[N+](=O)[O-] |
| CHK-A016 | 3JVS | CC(C)(C)c1ccccc1[N+](=O)[O-] |
| CHK-A017 | 3JVS | O=C(NNC(=O)c1cccc([N+](=O)[O-])c1)Nc1cccc2ccccc12 |
| CHK-A018 | 3JVS | O=CNNC(=O)Nc1cccc2ccccc12 |
| CHK-O000 | 3NLB | Cc1cc(-c2nc3cc(C(C)C)ccc3[nH]2)[nH]n1 |
| CHK-O001 | 3NLB | Cc1n[nH]c(-c2nc3cc(C(C)C)ccc3[nH]2)c1C(N)=O |
| CHK-O002 | 3NLB | Cc1n[nH]c(-c2nc3cc(C(C)C)ccc3[nH]2)c1C=O |
| CHK-O003 | 3NLB | Cc1n[nH]c(-c2nc3ccccc3[nH]2)c1C(=O)NC1CCN(C)CC1 |
| CHK-O004 | 3NLB | Cc1n[nH]cc1C(=O)NC1CCN(C)CC1 |
| CHK-O005 | 3OT3 | Cn1cc(-c2cnn3c(N)c(Br)cnc23)cn1 |
| CHK-O006 | 3OT3 | Nc1c(Br)c(C2CCCC(N)C2)nc2ccnn12 |
| CHK-O007 | 4HYI | NC(=O)c1csc(-n2ncc3ccccc32)n1 |
| CHK-O008 | 4HYI | O=C(Nc1ccccc1)c1csc(-n2ncc3ccccc32)n1 |
| CHK-O009 | 4HYI | O=C(Nc1ccccc1N1CCNCC1)c1cscn1 |
| CHK-O010 | 4HYI | O=CNc1ccccc1N1CCNCC1 |
| CHK-O011 | 4HYI | O=Cc1csc(-n2ncc3ccccc32)n1 |
| CHK-O012 | 4QYF | Nc1ncc(-c2cccc(O)c2)nc1-c1ccccc1 |
| CHK-O013 | 4QYF | Nc1nccnc1-c1ccc(C(=O)O)cc1 |
| CHK-O014 | 8SIV | CC1(N2CCC(c3cc4cc(N)ncc4cc3Cl)CC2)COCC1O |
| CHK-O015 | 8SIV | CC1(N2CCC(c3cc4cc(NC=O)ncc4cc3Cl)CC2)COCC1O |
| CHK-O016 | 8SIV | CC1(N2CCC(c3cc4ccncc4cc3Cl)CC2)COCC1O |
| CHK-O017 | 8SIV | O=C(Nc1cc2cc(C3CCNCC3)c(Cl)cc2cn1)C1CC1 |
| CHK-O018 | 8SIV | O=C(Nc1cc2ccc(Cl)cc2cn1)C1CC1 |
| CSK-A000 | 5MMF | CCC[NH2+]Cc1cccc(Cl)c1 |
| CSK-A001 | 5MMF | Cc1ccc(-c2ccccc2)c(Cl)c1 |
| CSK-A002 | 5MMF | Clc1ccccc1-c1ccccc1 |
| CSK-A003 | 5MMF | [NH3+]Cc1ccc(-c2ccccc2)c(Cl)c1 |
| CSK-A004 | 5MOD | CC(C)Oc1ccccc1Cl |
| CSK-A005 | 5MOD | NCc1ccc(O)c(Cl)c1 |
| CSK-A006 | 5OSU | COc1ccc(C(F)(F)F)cc1 |
| CSK-A007 | 5OSU | COc1ccc(C(F)(F)F)cc1-c1ccccc1Cl |
| CSK-A008 | 5OSU | COc1ccccc1-c1ccc(CN)cc1Cl |
| CSK-A009 | 5OSU | NCc1ccc(-c2cccc(C(F)(F)F)c2)c(Cl)c1 |
| CSK-A010 | 6GIH | Clc1cccc(-c2cccc3[nH]ccc23)c1 |
| CSK-O000 | 3MB7 | O=C(O)c1cc2c(ccc3ccc4occc4c32)o1 |
| CSK-O001 | 3R0T | FC(F)(F)c1cccc(Nc2nc3ccccc3c3cnc(NC4CC4)nc23)c1 |

Table S1 – continued from previous page

| Fragment ID | PDB | SMILES |
|---|---|---|
| CSK-O002 | 3R0T | Nc1cccc(C(F)(F)F)c1 |
| CSK-O003 | 3R0T | Nc1nc2cc(C(=O)O)ccc2c2cnc(NC3CC3)nc12 |
| CSK-O004 | 3R0T | Nc1ncc2c(n1)c(Nc1cccc(C(F)(F)F)c1)nc1cc(C(=O)O)ccc12 |
| CSK-O005 | 3R0T | O=C(O)c1ccc2c(c1)nc(Nc1cccc(C(F)(F)F)c1)c1ncncc12 |
| CSK-O006 | 3R0T | O=C(O)c1ccc2c(c1)nc(Nc1ccccc1)c1nc(NC3CC3)ncc12 |
| CSK-O007 | 3R0T | O=C(O)c1ccc2c(c1)ncc1nc(NC3CC3)ncc12 |
| CSK-O008 | 5M4F | O=c1c(O)c(-c2ccccc2)oc2c(Cl)cc(Cl)cc12 |
| CSK-O009 | 5M4F | O=c1c(O)coc2c(Cl)cc(Cl)cc12 |
| CSK-O010 | 6YUM | CC(C)(C)OC(=O)N(CCO)c1ccn2ncc(-c3ccc(C(=O)O)c(O)c3)c2n1 |
| CSK-O011 | 6YUM | CC(C)(C)OC(=O)N(CCOCCO)c1ccn2ncc(-c3cccc(O)c3)c2n1 |
| CSK-O012 | 6YUM | CC(C)(C)OC(=O)N(CCOCCO)c1ccn2nccc2n1 |
| CSK-O013 | 6YUM | CC(C)(C)OC(=O)NCCOCCO |
| CSK-O014 | 6YUM | CC(C)(C)OC(=O)Nc1ccn2ncc(-c3ccc(C(=O)O)c(O)c3)c2n1 |
| CSK-O015 | 6YUM | CCN(C(=O)OC(C)(C)C)c1ccn2ncc(-c3ccc(C(=O)O)c(O)c3)c2n1 |
| CSK-O016 | 6YUM | O=C(O)c1ccc(-c2cnn3ccc(N(CCOCCO)C(=O)O)nc23)cc1O |
| CSK-O017 | 6YUM | O=C(O)c1ccc(-c2cnn3ccc(NCCOCCO)nc23)cc1O |
| CSK-O018 | 6YUM | O=C(O)c1ccc(-c2cnn3cccnc23)cc1O |
| CSK-O019 | 6YUM | O=CN(CCOCCO)c1ccn2ncc(-c3ccc(C(=O)O)c(O)c3)c2n1 |
| MAP-A000 | 3NEW | O=c1cc(C(F)(F)F)c2c(-c3ccccc3)n[nH]c2[nH]1 |
| MAP-A001 | 3NEW | O=c1cc(C(F)(F)F)c2cn[nH]c2[nH]1 |
| MAP-A002 | 3NEW | O=c1ccc2c(-c3cccc(C(F)(F)F)c3)n[nH]c2[nH]1 |
| MAP-A003 | 5N63 | CNc1nc(-c2ccccc2)nc2cc(N)ccc12 |
| MAP-A004 | 5N63 | Nc1ccc2c(N)nc(-c3ccccc3)nc2c1 |
| MAP-A005 | 5N63 | Nc1ccc2c(NCc3ccc(F)cc3)ncnc2c1 |
| MAP-A006 | 5N63 | Nc1ccc2cnc(-c3ccccc3)nc2c1 |
| MAP-A007 | 5N64 | Nc1ccc2c(NCc3cccs3)ncnc2c1 |
| MAP-A008 | 5N67 | CC(=O)N1CCN(c2ccc(-c3nc(N)c4ccc(N)cc4n3)cc2)CC1 |
| MAP-A009 | 5N67 | CC(=O)N1CCN(c2ccc(-c3ncc4ccc(N)cc4n3)cc2)CC1 |
| MAP-A010 | 5N67 | CC(=O)N1CCN(c2ccccc2)CC1 |
| MAP-A011 | 5N67 | CCNc1nc(-c2ccc(N3CCN(C(C)=O)CC3)cc2)nc2cc(N)ccc12 |
| MAP-A012 | 5N67 | Nc1ccc2c(NCCc3ccccc3)nc(-c3ccc(N4CCNCC4)cc3)nc2c1 |
| MAP-A013 | 5N67 | Nc1ccc2c(NCCc3ccccc3)nc(-c3ccccc3)nc2c1 |
| MAP-A014 | 5N67 | Nc1ccc2c(NCCc3ccccc3)ncnc2c1 |
| MAP-A015 | 5N68 | CCNc1nc(-c2ccc(N3CCOCC3)cc2)nc2cc(N)ccc12 |
| MAP-A016 | 5N68 | Nc1ccc2c(N)nc(-c3ccc(N4CCOCC4)cc3)nc2c1 |
| MAP-A017 | 5N68 | Nc1ccc2cnc(-c3ccc(N4CCOCC4)cc3)nc2c1 |
| MAP-A018 | 5N68 | c1ccc(N2CCOCC2)cc1 |
| MAP-A019 | 8X3M | COc1cc2nc(N)sc2cc1OC |
| MAP-A020 | 8X3M | COc1cc2nc(NC(=O)c3csc(C)n3)sc2cc1OC |
| MAP-A021 | 8X3M | COc1cc2nc(NC(=O)c3csc(CO)n3)sc2cc1OC |
| MAP-A022 | 8X3M | COc1cc2nc(NC(=O)c3cscn3)sc2cc1OC |
| MAP-A023 | 8X3M | COc1cc2nc(NC=O)sc2cc1OC |
| MAP-A024 | 8X3M | COc1cc2ncsc2cc1OC |
| MAP-A025 | 8X3M | COc1ccc2nc(NC(=O)c3csc(COc4ccc(F)cc4)n3)sc2c1 |
| MAP-A026 | 8X3M | COc1ccc2sc(NC(=O)c3csc(COc4ccc(F)cc4)n3)nc2c1 |
| MAP-A027 | 8X3M | Fc1ccc(OCc2nccs2)cc1 |
| MAP-A028 | 8X3M | NC(=O)c1csc(COc2ccc(F)cc2)n1 |
| MAP-A029 | 8X3M | O=Cc1csc(COc2ccc(F)cc2)n1 |
| MAP-A030 | 8YD9 | COc1ccc(C2NC(N)=Nc3nc4cc5c(cc4n32)OCCCO5)cc1 |
| MAP-A031 | 8YD9 | NC1=Nc2nc3cc4c(cc3n2C(c2ccc(O)cc2)N1)OCCCO4 |
| MAP-A032 | 8YD9 | NC1=Nc2nc3cc4c(cc3n2C(c2ccccc2)N1)OCCCO4 |
| MAP-O000 | 1OUK | CC(Nc1nccc(-c2c(-c3cccc(C(F)(F)F)c3)ncn2C)n1)c1ccccc1 |
| MAP-O001 | 1OUK | CC(Nc1nccc(-c2c(-c3ccccc3)nc(C3CCNCC3)n2C)n1)c1ccccc1 |
| MAP-O002 | 1OUK | CC(Nc1nccc(-c2cnc(C3CCNCC3)n2C)n1)c1ccccc1 |
| MAP-O003 | 1OUK | CC(Nc1ncccn1)c1ccccc1 |
| MAP-O004 | 1OUK | CCNc1nccc(-c2c(-c3cccc(C(F)(F)F)c3)nc(C3CCNCC3)n2C)n1 |
| MAP-O005 | 1OUK | Cn1c(C2CCNCC2)nc(-c2cccc(C(F)(F)F)c2)c1-c1ccnc(N)n1 |
| MAP-O006 | 1OUK | Cn1c(C2CCNCC2)nc(-c2cccc(C(F)(F)F)c2)c1-c1ccncn1 |
| MAP-O007 | 1OUK | Cn1cc(-c2cccc(C(F)(F)F)c2)nc1C1CCNCC1 |
| MAP-O008 | 2YIX | CC(C)c1nnc2ccc(Sc3ccccc3)cn12 |
| MAP-O009 | 2YIX | CC(C)c1nnc2ccc(Sc3ccccc3CN)cn12 |
| MAP-O010 | 2YIX | CC(C)c1nnc2ccc(Sc3ccccc3CNC(N)=O)cn12 |

Table S1 – continued from previous page

| Fragment ID | PDB | SMILES |
|---|---|---|
| MAP-O011 | 2YIX | CC(C)c1nnc2ccc(Sc3ccccc3CNC=O)cn12 |
| MAP-O012 | 2YIX | CCNC(=O)NCc1ccccc1S |
| MAP-O013 | 2YIX | CCNC(=O)NCc1ccccc1Sc1ccc2nncn2c1 |
| MAP-O014 | 2YIX | Cc1ccccc1Sc1ccc2nnc(C(C)C)n2c1 |
| MAP-O015 | 3GFE | Cc1ccc(C(=O)NC2CC2)cc1Nc1cc(=O)n(C)c2[nH]ncc12 |
| MAP-O016 | 3GFE | Cc1ccc(C(N)=O)cc1Nc1cc(=O)n(C)c2c1cnn2-c1ccc(F)cc1F |
| MAP-O017 | 3GFE | Cc1ccc(C=O)cc1Nc1cc(=O)n(C)c2c1cnn2-c1ccc(F)cc1F |
| MAP-O018 | 3GFE | Cc1ccccc1Nc1cc(=O)n(C)c2c1cnn2-c1ccc(F)cc1F |
| MAP-O019 | 3GFE | Cn1c(=O)cc(N)c2cnn(-c3ccc(F)cc3F)c21 |
| MAP-O020 | 3GFE | Cn1c(=O)ccc2cnn(-c3ccc(F)cc3F)c21 |
| MAP-O021 | 3U8W | CCn1c(=O)c(-c2cc(C(=O)NC3CC3)ccc2C)cc2nncn21 |
| MAP-O022 | 3U8W | CCn1c(=O)c(-c2cc(C(N)=O)ccc2C)cc2nnc(-c3c(F)cccc3Cl)n21 |
| MAP-O023 | 3U8W | CCn1c(=O)c(-c2cc(C=O)ccc2C)cc2nnc(-c3c(F)cccc3Cl)n21 |
| MAP-O024 | 3U8W | CCn1c(=O)c(-c2ccccc2C)cc2nnc(-c3c(F)cccc3Cl)n21 |
| MAP-O025 | 3U8W | CCn1c(=O)ccc2nnc(-c3c(F)cccc3Cl)n21 |
| MAP-O026 | 3U8W | Cc1ccc(C(=O)NC2CC2)cc1-c1cc2nnc(-c3c(F)cccc3Cl)n2[nH]c1=O |
| MAP-O027 | 4KIN | Cc1ccc(C(=O)NC2CC2)cc1NC(=O)c1cccs1 |
| MAP-O028 | 4KIN | Cc1ccc(C(=O)NC2CC2)cc1NC=O |
| MAP-O029 | 4KIN | Cc1ccc(C(N)=O)cc1NC(=O)c1ccc(-c2ccccc2Cl)s1 |
| MAP-O030 | 4KIN | Cc1ccc(C=O)cc1NC(=O)c1ccc(-c2ccccc2Cl)s1 |
| MAP-O031 | 4KIN | Cc1ccccc1NC(=O)c1ccc(-c2ccccc2Cl)s1 |
| MAP-O032 | 4KIN | Clc1ccccc1-c1cccs1 |
| MAP-O033 | 4KIN | NC(=O)c1ccc(-c2ccccc2Cl)s1 |
| MAP-O034 | 4KIN | O=Cc1ccc(-c2ccccc2Cl)s1 |

**Table S2:** Spearman $\rho$ correlations between molecular descriptors and orthosteric memorization rate in the Co-FFF non-orthosteric exploration screen, without and with ADP blocker ($n = 232$ for all descriptors except Tanimoto, $n = 116$). Orthosteric memorization rate is the per-fragment fraction of cofold samples localized to the orthosteric site (fragment center of mass to the nearest heavy-atom distance $\leq 5$ Å), pooled across five kinases and ten diffusion samples per fragment.

| **Descriptor** | **No ADP** | | **ADP blocker** | |
|---|---|---|---|---|
| | $\rho$ | $p$ | $\rho$ | $p$ |
| Aromatic rings | +0.427 | < 0.0001 | +0.079 | 0.230 |
| HBA | +0.386 | < 0.0001 | +0.170 | 0.0093 |
| N count | +0.330 | < 0.0001 | +0.081 | 0.220 |
| TPSA | +0.310 | < 0.0001 | +0.241 | 0.0002 |
| N rings | +0.273 | < 0.0001 | −0.001 | 0.984 |
| HBD | +0.214 | 0.0010 | +0.340 | < 0.0001 |
| Halogen count | −0.206 | 0.0016 | −0.102 | 0.122 |
| Fsp3 | −0.202 | 0.0020 | −0.010 | 0.876 |
| Tanimoto | +0.167 | 0.0737 | −0.052 | 0.583 |
| Heavy atoms | +0.143 | 0.0291 | +0.050 | 0.445 |
| MW | +0.117 | 0.0754 | +0.069 | 0.294 |
| LogP | −0.043 | 0.510 | −0.174 | 0.0079 |
| Rotatable bonds | +0.003 | 0.969 | +0.157 | 0.0165 |

**Table S3:** Non-orthosteric exploration rate (%) for all five kinase targets under three blocking conditions: no blocker, Type I inhibitor, and ADP. Values are mean non-orthosteric exploration rate across all 232 fragments per target (100 − orthosteric localization rate). The Type I blocker for each kinase was extracted from the indicated PDB structure using the corresponding CCD ligand code.

| Kinase | Type I PDB | CCD | No blocker | Type I blocker | ADP blocker |
|---|---|---|---|---|---|
| AKT2 | 2JDO | I5S | 7.2% | 92.0% | 97.7% |
| CDK2 | 2UUE | MTZ | 2.2% | 98.9% | 85.3% |
| CHEK1 | 2YEX | YEX | 14.4% | 84.8% | 98.3% |
| CSNK2A1 | 3WAR | NIO | 20.8% | 83.3% | 91.1% |
| MAPK14 | 3S3I | CQ0 | 13.8% | 95.9% | 52.5% |
| Mean | | | 11.6% | 91.0% | 85.0% |

**Table S4:** Boltz allosteric pose vs. crystallographic reference pose $\Delta G$ (kcal/mol) for the 30 fragment–kinase complexes shown in Figure 4. Where the same fragment (shared ID) was tested against more than one parent structure, the source PDB is given parenthetically. Orthosteric pose $\Delta G$ is additionally reported for the 10 fragments that showed clean ADP-dependent redirection (i.e., 100% orthosteric occupancy in the unblocked (no-ADP) condition, with the fragment relocating to the allosteric pocket only upon ADP blocking).

| Fragment ID | Crystal $\Delta G$ | Allo $\Delta G$ | Ortho $\Delta G$ |
|---|---|---|---|
| AKT-A008 | $1.07 \pm 0.18$ | $-3.79 \pm 0.17$ | $-1.86 \pm 0.15$ |
| AKT-A009 | $-14.90 \pm 0.41$ | $-12.62 \pm 0.40$ | — |
| AKT-A010 | $-11.34 \pm 0.33$ | $-7.14 \pm 0.26$ | — |
| CDK-A004 | $-1.70 \pm 0.10$ | $-0.74 \pm 0.09$ | $1.22 \pm 0.15$ |
| CHK-A003 | $0.10 \pm 0.37$ | $-0.28 \pm 0.35$ | $-4.67 \pm 0.42$ |
| CHK-A006 | $-2.57 \pm 0.11$ | $-3.29 \pm 0.11$ | $-1.23 \pm 0.17$ |
| CHK-A015 | $-5.11 \pm 0.15$ | $-5.07 \pm 0.15$ | — |
| CHK-A017 | $-5.31 \pm 0.31$ | $-5.70 \pm 0.26$ | — |
| CSK-A000 | $-4.82 \pm 0.19$ | $-6.51 \pm 0.16$ | — |
| CSK-A001 | $-6.86 \pm 0.10$ | $-7.32 \pm 0.09$ | — |
| CSK-A002 | $-7.01 \pm 0.08$ | $-7.21 \pm 0.08$ | — |
| CSK-A003 | $-8.08 \pm 0.15$ | $-9.06 \pm 0.12$ | — |
| CSK-A004 | $-4.67 \pm 0.10$ | $-6.83 \pm 0.08$ | — |
| CSK-A005 | $-1.37 \pm 0.15$ | $-1.38 \pm 0.12$ | — |
| CSK-A006 | $-6.44 \pm 0.08$ | $-7.13 \pm 0.06$ | — |
| CSK-A007 | $-11.45 \pm 0.10$ | $-11.57 \pm 0.11$ | — |
| CSK-A008 | $-10.11 \pm 0.13$ | $-10.44 \pm 0.13$ | — |
| CSK-A009 | $-8.98 \pm 0.12$ | $-9.53 \pm 0.12$ | — |
| MAP-A000 | $-5.77 \pm 0.20$ | $-7.32 \pm 0.22$ | $-3.49 \pm 0.17$ |
| MAP-A003 (5N63) | $-6.04 \pm 0.23$ | $-4.38 \pm 0.22$ | $-5.43 \pm 0.19$ |
| MAP-A003 (5N64) | $-0.86 \pm 0.30$ | $-1.29 \pm 0.27$ | — |
| MAP-A004 (5N63) | $-4.80 \pm 0.27$ | $-4.06 \pm 0.22$ | — |
| MAP-A004 (5N64) | $-3.51 \pm 0.29$ | $-5.89 \pm 0.22$ | $-5.19 \pm 0.15$ |
| MAP-A005 | $-6.43 \pm 0.25$ | $-5.80 \pm 0.24$ | $-7.65 \pm 0.17$ |
| MAP-A006 (5N63) | $-4.10 \pm 0.21$ | $-3.20 \pm 0.23$ | — |
| MAP-A006 (5N64) | $-0.63 \pm 0.29$ | $-0.68 \pm 0.22$ | — |
| MAP-A007 | $-2.80 \pm 0.21$ | $-4.19 \pm 0.23$ | $-6.23 \pm 0.15$ |
| MAP-A010 | $1.02 \pm 0.13$ | $0.35 \pm 0.16$ | — |
| MAP-A014 | $-7.23 \pm 0.23$ | $-4.00 \pm 0.30$ | $-7.50 \pm 0.15$ |
| MAP-A017 | $-6.80 \pm 0.26$ | $-9.91 \pm 0.24$ | — |

**Table S5:** ABFE selectivity results for INX-315 and BRICS-derived fragments against CDK2 and CDK6 in orthosteric poses without ADP blocker, phosphorylated activation state. All values in kcal/mol, $n = 2$ Boltz models per condition.

| Ligand | SMILES | HA | $\Delta G_{\text{CDK2}}$ | $\Delta G_{\text{CDK6}}$ |
|---|---|---|---|---|
| INX-315 | `C1CCC2(CC1)C(=O)NNC3=CC4=CN=C(N=C4N23)C5=CC(=C5)S(=O)(=O)N` | 30 | −9.03 | −7.85 |
| Frag 1 | `Nc1ccc(S(N)(=O)=O)cc1` | 11 | −0.87 | +0.52 |
| Frag 2 | `Nc1ncc2cn(c2n1)C1(CCCC1)C(=O)NN3` | 20 | −4.92 | −2.38 |
| Frag 3 | `NS(=O)(=O)c1ccccc1` | 10 | +2.47 | −1.07 |

**Table S6:** Enamine screen: all 33 favorable ($\Delta G \leq -1$ kcal/mol) allosteric/cryptic poses, alongside the orthosteric-pose $\Delta G$ where available. Nonbinding indicates a positive binding free energy or an unstable simulation. Blank orthosteric entries indicate that no orthosteric pose was obtained in the no ADP screen.

| Kinase | Catalog ID | Pocket | Allo/Cryptic $\Delta G$ | Ortho $\Delta G$ |
|---|---|---|---|---|
| AKT2 | Z431983770 | cryptic | $-3.77 \pm 0.17$ | $-0.45 \pm 0.15$ |
| AKT2 | Z1258538520 | cryptic | $-1.34 \pm 0.17$ | $-3.03 \pm 0.20$ |
| AKT2 | Z997587036 | cryptic | $-1.12 \pm 0.14$ | $-1.94 \pm 0.11$ |
| AKT2 | Z227266274 | cryptic | $-1.07 \pm 0.15$ | $-4.17 \pm 0.13$ |
| CDK2 | Z1726128749 | allo | $-5.43 \pm 12.48$ | nonbinding |
| CDK2 | Z1198223612 | allo | $-2.05 \pm 0.11$ | $-4.22 \pm 0.15$ |
| CDK2 | Z1192359567 | allo | $-1.24 \pm 0.12$ | nonbinding |
| CDK2 | Z57683752 | allo | $-1.15 \pm 0.13$ | nonbinding |
| CDK2 | Z734660320 | cryptic | $-1.33 \pm 0.16$ | nonbinding |
| CHEK1 | Z1270107479 | allo | $-4.66 \pm 0.10$ | nonbinding |
| CHEK1 | Z1262562584 | allo | $-4.02 \pm 0.08$ | nonbinding |
| CHEK1 | Z1509503466 | allo | $-2.97 \pm 0.11$ | nonbinding |
| CHEK1 | Z381402572 | allo | $-2.04 \pm 0.10$ | nonbinding |
| CHEK1 | Z1198223612 | cryptic | $-2.73 \pm 0.13$ | nonbinding |
| CHEK1 | Z1098229000 | cryptic | $-2.66 \pm 0.12$ | $-1.40 \pm 0.13$ |
| CSNK2A1 | Z1154602987 | allo | $-4.49 \pm 0.13$ | nonbinding |
| CSNK2A1 | Z196118762 | allo | $-3.74 \pm 0.11$ | $-4.05 \pm 0.15$ |
| CSNK2A1 | Z1098229000 | allo | $-3.68 \pm 0.10$ | $-3.59 \pm 0.13$ |
| CSNK2A1 | Z422382038 | allo | $-2.11 \pm 0.14$ | nonbinding |
| CSNK2A1 | Z975853462 | allo | $-2.00 \pm 0.11$ | $-7.17 \pm 0.16$ |
| CSNK2A1 | Z1198313078 | allo | $-1.36 \pm 0.10$ | $-3.57 \pm 0.12$ |
| CSNK2A1 | Z362847974 | allo | $-1.11 \pm 0.10$ | nonbinding |
| MAPK14 | Z1255396793 | allo | $-2.56 \pm 0.09$ | — |
| MAPK14 | Z1255448870 | allo | $-2.10 \pm 0.10$ | nonbinding |
| MAPK14 | Z2235790608 | allo | $-1.41 \pm 0.16$ | $-2.98 \pm 0.12$ |
| MAPK14 | Z1255453358 | allo | $-1.09 \pm 0.11$ | nonbinding |
| MAPK14 | Z1551916919 | allo | $-1.02 \pm 0.13$ | — |
| MAPK14 | Z425721880 | cryptic | $-1.82 \pm 0.15$ | nonbinding |
| MAPK14 | Z1098324193 | cryptic | $-1.80 \pm 0.15$ | $-2.85 \pm 0.18$ |
| MAPK14 | Z1171972190 | cryptic | $-1.17 \pm 0.13$ | nonbinding |
| MAPK14 | Z1198237136 | cryptic | $-1.14 \pm 0.15$ | nonbinding |
| MAPK14 | Z1477409275 | cryptic | $-1.13 \pm 0.11$ | nonbinding |
| MAPK14 | Z1118527729 | cryptic | $-1.12 \pm 0.13$ | — |

### 2 Supplementary Figures

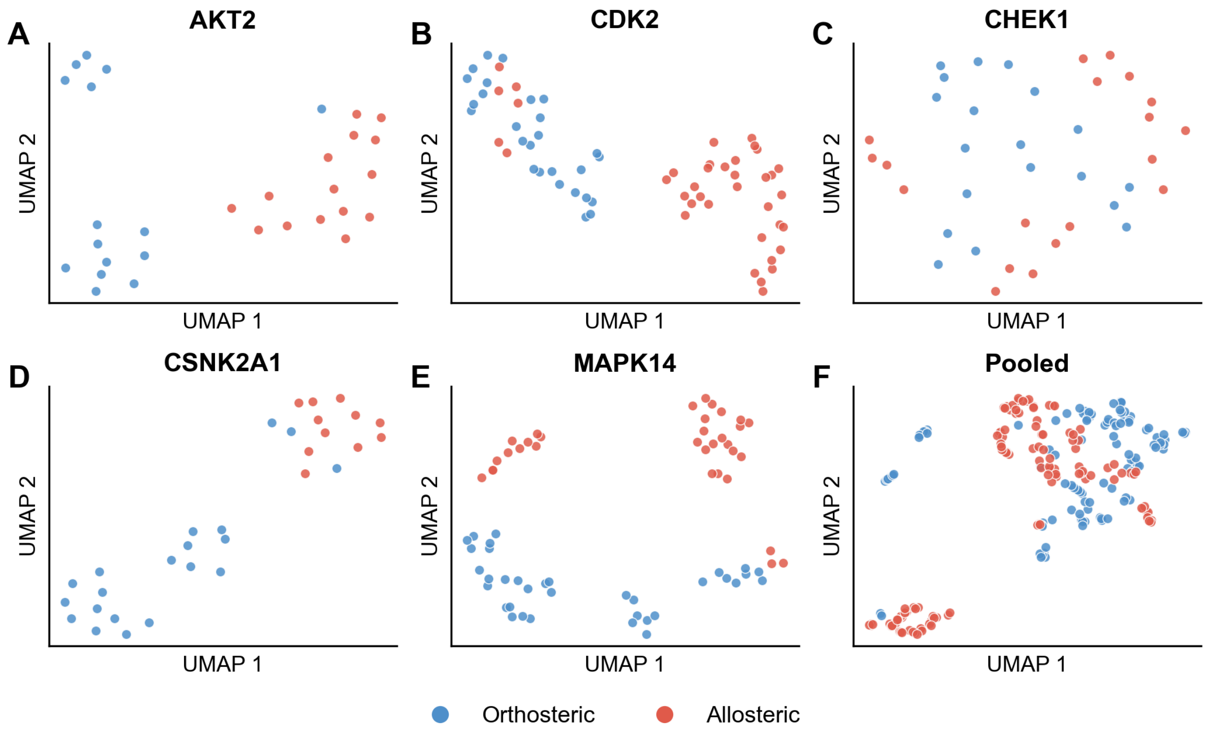


**Figure S1: UMAP embedding of orthosteric and allosteric single-cut BRICS fragments across five kinases.** (A–E) Per-kinase panels: AKT2, CDK2, CHEK1, CSNK2A1, and MAPK14, with UMAP fit independently on each kinase's fragments. (F) Pooled embedding across all five kinases, fit on 116 unique orthosteric and 116 unique allosteric fragments (deduplicated by SMILES). Fingerprints: RDKit Morgan (radius 2, 2048 bits). Dimensionality reduction: UMAP with Jaccard metric, n_neighbors = 15, min_dist = 0.1, random_state = 42. Because panels A–E are fit independently, UMAP1/UMAP2 coordinates are not comparable across panels; only within-panel clustering (orthosteric vs. allosteric separation) is meaningful.

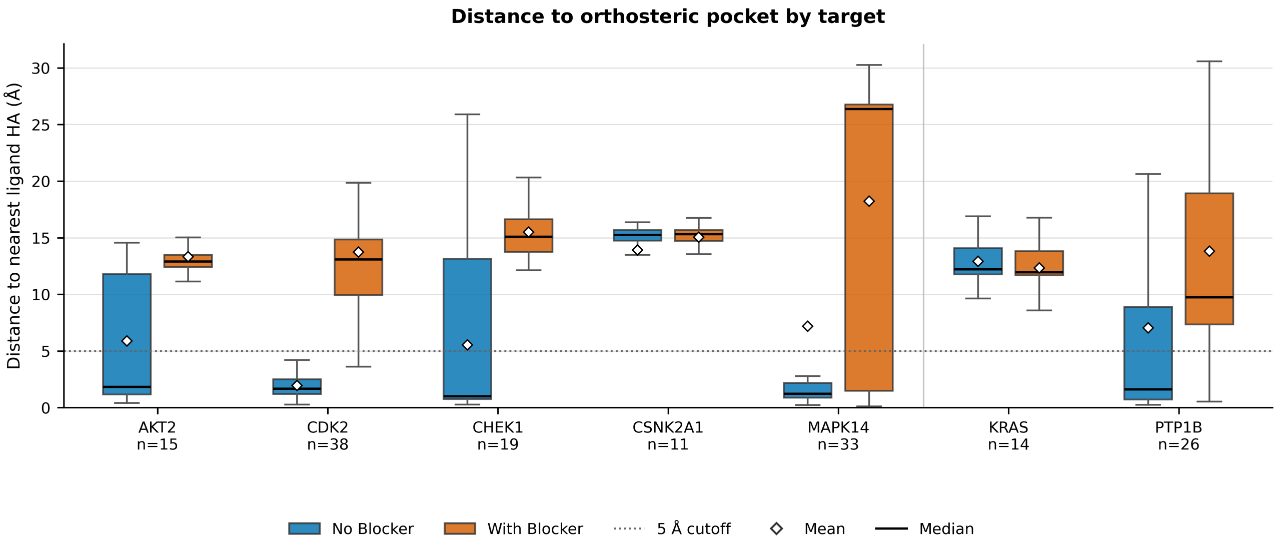


**Figure S2: Distance of predicted fragment poses to the orthosteric pocket across targets with and without an orthosteric blocker.** Distance is from the predicted fragment heavy-atom center of mass to the nearest heavy atom of the orthosteric reference ligand. The dotted line marks the 5 Å localization cutoff used to classify a fragment sample as orthosterically localized. Sample sizes ($n$) are the number of fragments per target. Kinases are shown left of the vertical divider; KRAS and PTP1B are non-kinase controls.

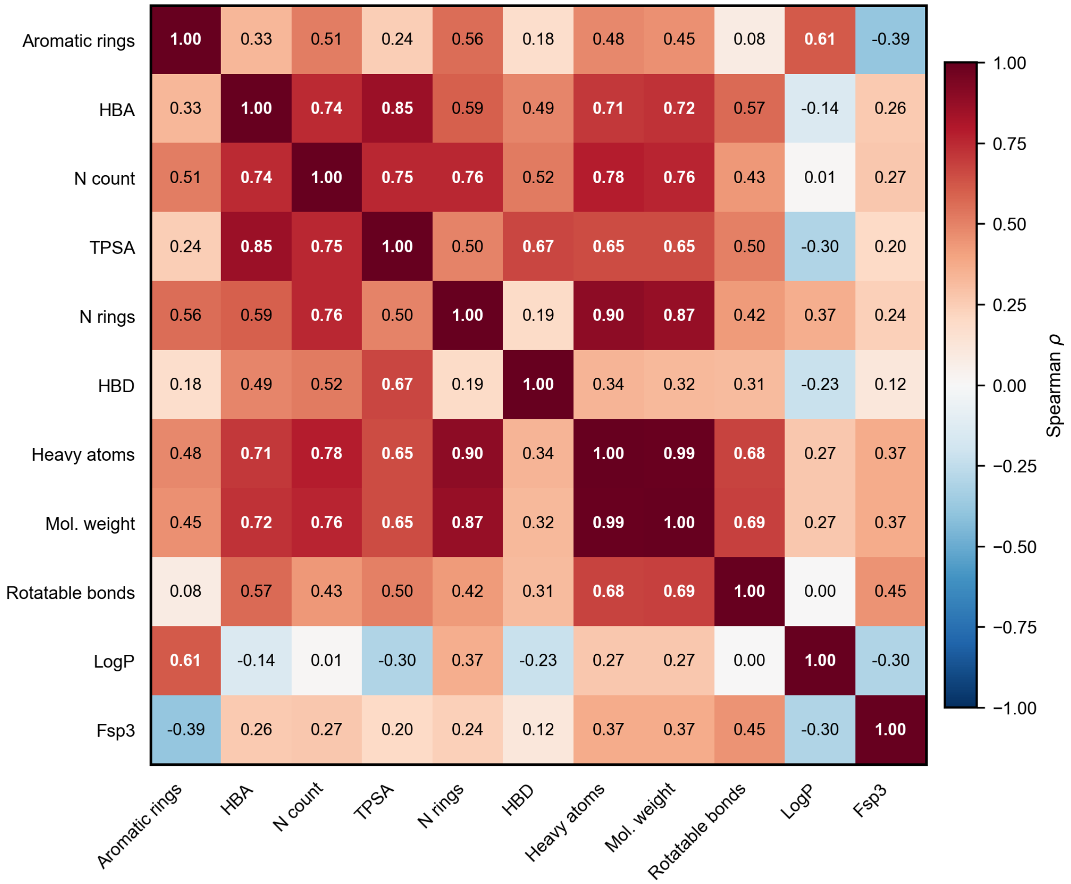


**Figure S3: Inter-descriptor correlation matrix for the fragment library.** Spearman $\rho$ values between all pairs of molecular descriptors ($n = 232$). HBA, N count, and TPSA form a tightly correlated cluster ($\rho \geq 0.74$), as do heavy atoms and MW ($\rho = 0.99$), indicating substantial redundancy among size- and polarity-related descriptors. Fsp3 and LogP are largely orthogonal to the remaining descriptors. Correlations are independent of ADP blocking condition.

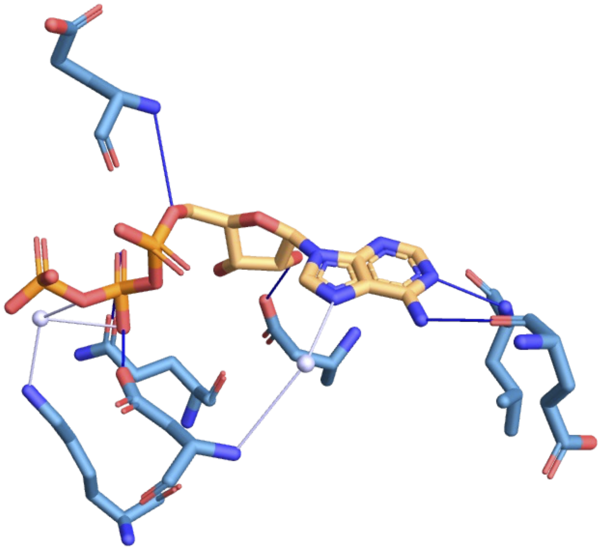

**Figure S4: ATP makes three nitrogen-mediated contacts in the CDK2 orthosteric pocket (PDB: 1FIN).** The adenine moiety of ATP (orange) forms hydrogen bonds (blue lines) through its three ring nitrogens with hinge region residues of CDK2 (teal). Magnesium ions are shown as white spheres coordinating the phosphate groups. Fragments with more aromatic rings, hydrogen bond acceptors, and nitrogen atoms are more likely to recapitulate this pharmacophore, partially driving orthosteric memorization in co-folding predictions.

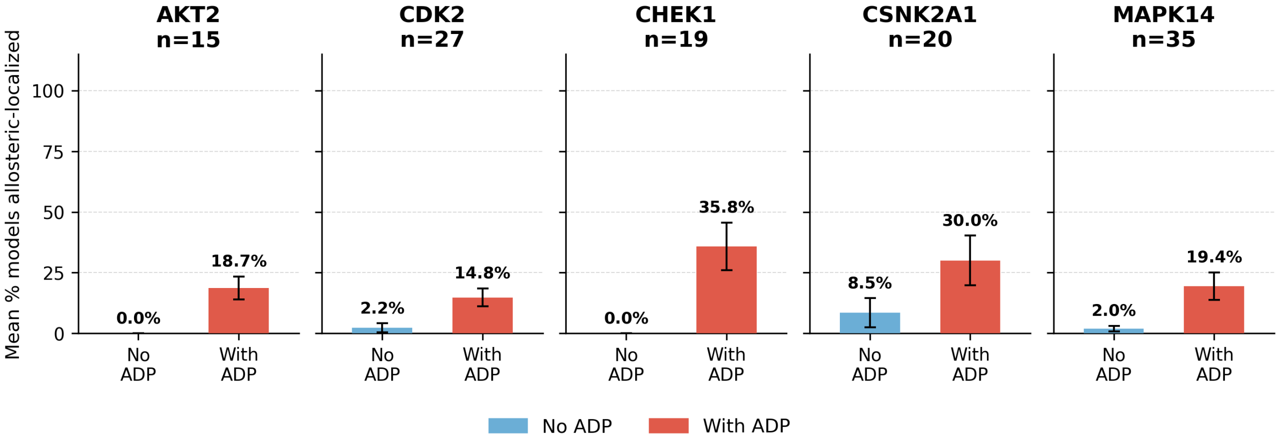


**Figure S5:** Allosteric-pocket localization of orthosteric-derived fragments, with and without ADP blocking.
